# The Accumulation of Progerin Underlies the Loss of Aortic Smooth Muscle Cells in Hutchinson-Gilford Progeria Syndrome

**DOI:** 10.1101/2024.10.29.620896

**Authors:** Paul H. Kim, Joonyoung R. Kim, Patrick J. Heizer, Hyesoo Jung, Yiping Tu, Ashley Presnell, Julia Scheithauer, Rachel G. Yu, Stephen G. Young, Loren G. Fong

**Author notes:** To whom correspondence should be addressed: Loren G. Fong, Ph.D. or Stephen G. Young, M.D., 695 Charles E. Young Dr. South, Los Angeles, CA 90095. E-mail addresses.

## Abstract

Hutchinson-Gilford progeria syndrome (HGPS) is a progeroid disorder characterized by multiple aging-like phenotypes, including disease in large arteries. HGPS is caused by an internally truncated prelamin A (progerin) that cannot undergo the ZMPSTE24-mediated processing step that converts farnesyl-prelamin A to mature lamin A; consequently, progerin retains a carboxyl-terminal farnesyl lipid anchor. In cultured cells, progerin and full-length farnesyl-prelamin A (produced in *Zmpste24*^−/–^ cells) form an abnormal nuclear lamin meshwork accompanied by nuclear membrane ruptures and cell death; however, these proteins differ in their capacity to cause arterial disease. In a mouse model of HGPS (*Lmna*^G609G^), progerin causes loss of aortic smooth muscle cells (SMCs) by ∼12 weeks of age. In contrast, farnesyl-prelamin A in *Zmpste24*^−/–^ mice does not cause SMC loss—even at 21 weeks of age. In young mice, aortic levels of farnesyl-prelamin A in *Zmpste24*^−/–^ mice and aortic levels of progerin in *Lmna*^G609G/+^ mice are the same. However, the levels of progerin and other A-type lamins increase with age in *Lmna*^G609G/+^ mice, whereas farnesyl-prelamin A and lamin C levels in *Zmpste24*^−/–^ mice remain stable. *Lmna* transcript levels are similar, implying that progerin influences nuclear lamin turnover. We identified a likely mechanism. In cultured SMCs, the phosphorylation of Ser-404 by AKT (which triggers prelamin A degradation) is reduced in progerin. In mice, AKT activity is significantly lower in *Lmna*^G609G/+^ aortas than in wild-type or *Zmpste24*^−/–^ aortas. Our studies identify that the accumulation of progerin in *Lmna*^G609G^ aortas underlies the hallmark arterial pathology in HGPS.

**One Sentence Summary:** The age-related accumulation of progerin in smooth muscle cells (SMCs) explains the loss of arterial SMCs in Hutchinson-Gilford progeria syndrome.

## Introduction

Hutchinson-Gilford progeria syndrome (HGPS) is a pediatric progeroid disorder caused by point mutations that optimize a cryptic splice-donor site in exon 11 of *LMNA* (the gene for prelamin A and lamin C), resulting in aberrant mRNA splicing and the production of a prelamin A transcript lacking the last 150 nucleotides of exon 11 (1, 2). That transcript results in the production of a mutant prelamin A, commonly called progerin, containing an internal deletion of 50 amino acids. Children with HGPS can appear normal at birth, but they soon develop disease phenotypes that resemble, at least superficially, the phenotypes occurring with physiologic aging (3–5). Children with HGPS typically die during their teenage years from heart attacks or strokes (due to occlusive lesions in the cerebral and coronary arteries).

The products of the *LMNA* gene, prelamin A (the precursor to mature lamin A) and lamin C, are produced by alternative splicing (6). The lamin C transcript contains exon 1–10 sequences; prelamin A contains exon 1–12 sequences. Lamin C is identical to prelamin A through the first 566 amino acids but contains six unique residues at its C terminus. Prelamin A contains 98 unique C-terminal amino acids and terminates with a *CaaX* motif, which triggers protein farnesylation, followed by three additional processing steps [endoproteolytic release of the last three residues of the *CaaX* motif by RCE1 (“*aaXing*”), methylation of the newly exposed farnesylcysteine by ICMT, and endoproteolytic release of the last 15 amino acids (including the C-terminal farnesylcysteine methyl ester) by ZMPSTE24 (7)]. The ZMPSTE24-mediated processing step results in the production of mature lamin A.

The internal 50-amino acid deletion in progerin leaves its *CaaX* motif intact (and therefore does not affect protein farnesylation, *aaXing*, or methylation) but it eliminates the sequences required for ZMPSTE24 processing (8). Thus, progerin is farnesylated and methylated but cannot be processed to mature lamin A. Progerin’s C-terminal farnesylcysteine methyl ester enhances interactions with the inner nuclear membrane and limits the mobility of progerin. Fluorescence recovery studies following photobleaching revealed that the mobility of progerin is similar to that of lamin B1 (which is farnesylated and methylated) but slower than mature lamin A (which lacks the farnesyl lipid anchor) (9).

Progerin is the culprit molecule in disease pathogenesis. When progerin is expressed in cultured cells, it triggers misshapen nuclei, DNA damage, and cell senescence; these same abnormalities are observed in fibroblasts from patients with HGPS (1, 9–12). When progerin is expressed in mice, it triggers multiple disease phenotypes, including alopecia, bone fractures, and progressive weight loss (13–15). In addition, progerin causes the loss of smooth muscle cells (SMCs) in large arteries (16, 17). Loss of medial SMCs has also been documented in autopsy studies of aortic tissue from patients with HGPS (18, 19). Studies in mouse models of HGPS have provided valuable insights into the mechanisms of the vascular disease (15, 20–25). Of note, the high progerin*–*low lamin B1 nuclear lamin profile in aortic SMCs (which weakens the functional integrity of the nuclear envelope) combined with the rhythmic stretching/relaxation occurring in the wall of the aorta, play important roles in the tissue-specific loss of cells in the aorta.

Progerin’s ability to elicit disease phenotypes is often attributed to the fact that progerin, unlike mature lamin A or lamin C, contains a C-terminal farnesyl lipid. In support of that view, a protein farnesyltransferase inhibitor (FTI) improves nuclear shape in progerin-expressing cells and ameliorates disease in knock-in and transgenic models of HGPS (11–13, 16, 26–28). Also, disease phenotypes are absent in knock-in mouse models expressing a nonfarnesylated version of progerin (due to a single amino acid deletion in progerin’s *CaaX* motif) (29). Consistent with those observations, progerin in cultured cells results in the formation of a morphologically abnormal nuclear lamin meshwork (30, 31), whereas the meshwork formed by nonfarnesylated progerin is morphologically normal (31). The contribution of the internal deletion to the biological properties of progerin (other than eliminating ZMPSTE24-dependent processing) is less clear.

Studies of *Zmpste24*^−/–^ mice have provided further insights into the pathogenesis of progeria (32). ZMPSTE24 deficiency abolishes the conversion of farnesyl-prelamin A to mature lamin A; consequently, tissues of *Zmpste24*^−/–^ mice contain lamin C and farnesyl-prelamin A but lack mature lamin A. In humans, ZMPSTE24 deficiency causes restrictive dermopathy, a neonatal-lethal progeroid syndrome characterized by markedly impaired growth, rigid skin, joint contractures, and bone fractures (33). In mice, ZMPSTE24 deficiency causes many of the same phenotypes observed in gene-targeted models of HGPS (*e.g.*, failure-to-thrive, bone fractures, loss of adipose tissue); those phenotypes are severe and invariably result in premature death (34, 35). However, in contrast to the gene-targeted HGPS mice, *Zmpste24*^−/–^ mice or *Lmna*^L648R/L648R^ knock-in mice (which produce full-length farnesyl-prelamin A due to the blockade of ZMPSTE24 processing) do not exhibit loss of medial SMCs in the aorta (15, 36).

Why the HGPS mouse models, but not *Zmpste24*^−/–^ mice, exhibit vascular pathology is unknown, but this difference suggests that progerin and full-length farnesyl-prelamin A—despite both having identical C-terminal modifications—could have very distinct effects in the aorta. In the current study, we investigated that possibility. We discovered that progerin, but not farnesyl-prelamin A, accumulates progressively in SMCs in the mouse aorta, and that difference underlies their distinct capacities to cause arterial disease.

## Results

### Loss of vascular SMCs in HGPS knock-in mice but not *Zmpste24*^−/–^ mice

We examined aortas from wild-type mice (*Lmna*^+/+^), *Zmpste24*^−/–^ mice, and a gene-targeted knock-in mouse model of HGPS (*Lmna*^G609G/G609G^). Cross sections of the proximal ascending thoracic aorta were stained with antibodies against α-smooth muscle actin [a marker of SMCs], CD31 (an endothelial cell marker), and collagen type VIII [a marker of HGPS-associated adventitial fibrous; (15)]. Confocal microscopy images of the inner curvature of the ascending aorta in *Lmna*^G6096G/G609G^ mice revealed reduced α-smooth muscle actin staining, reduced numbers of SMCs, and increased collagen type VIII staining of the adventitia (**Fig. 1a** and **Supplementary Fig. 1a**). In contrast, aortas of *Zmpste24*^−/–^ mice were normal, indistinguishable from those in *Lmna*^+/+^ mice. Consistent with those findings, nuclear membrane ruptures (identified by escape of a nuclear-targeted tdTomato into the cytoplasm) were frequent in the medial Ss of *Lmna*^G6096G/G609G^ aortas but absent in *Zmpste24*^−/–^ aortas (**Fig. 1b** and **Supplementary Fig. 1b**). As expected, farnesyl-prelamin A was detected in aortas of *Zmpste24*^−/–^ mice but not in *Lmna*^+/+^ or *Lmna*^G6096G/G609G^ mice (**Fig. 1b** and **Supplementary Fig. 1b**). Numbers of vascular SMCs in the medial layer of the aorta were the same in *Zmpste24*^−/–^ mice and in wild-type mice (**Figs. 1c–d** and **Supplementary Fig. 2**).

**Fig. 1.**
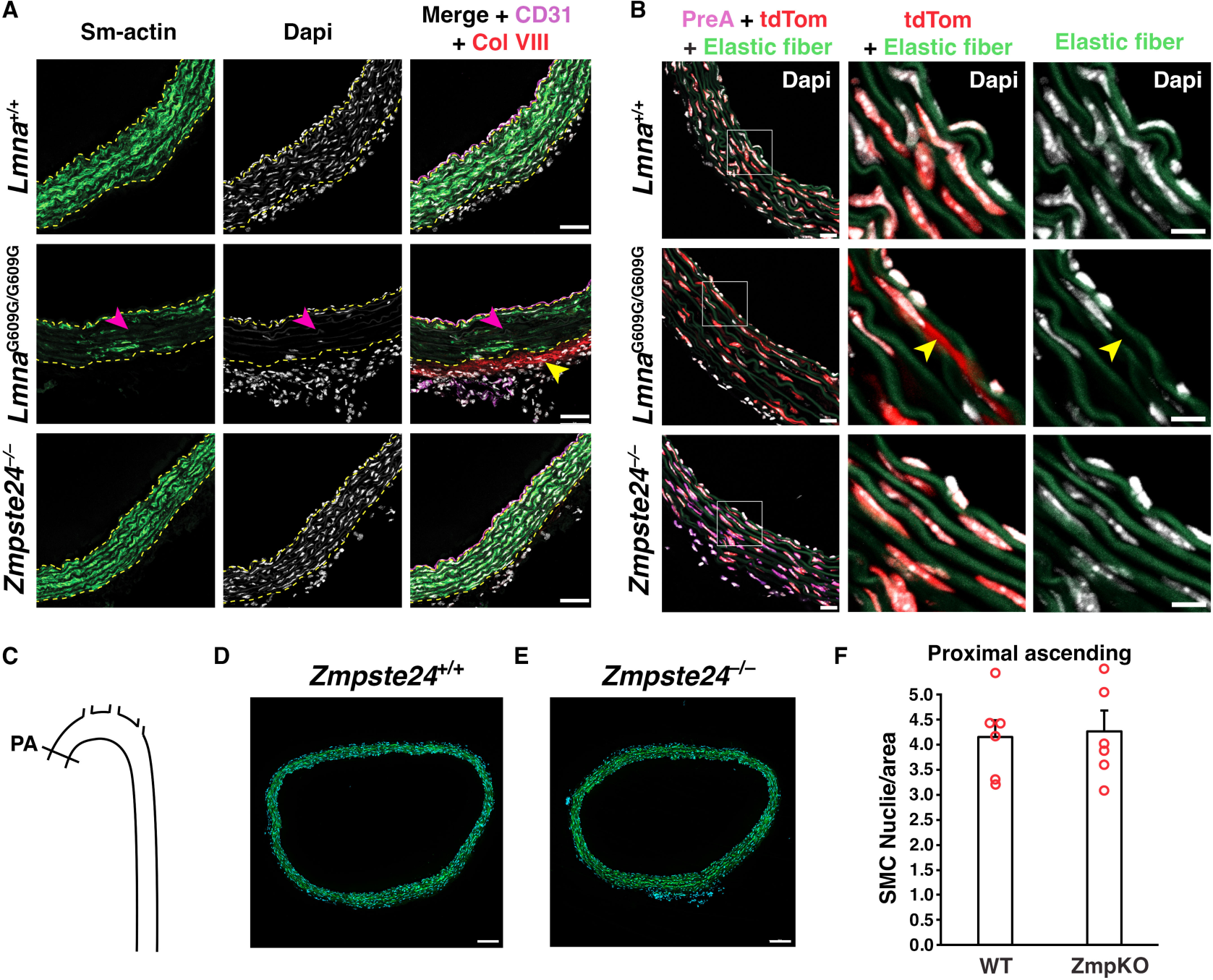
Loss of aortic smooth muscle cells (SMCs) occurs in HGPS knock-in mice but not *Zmpste24*^−/–^ mice. **A**. Confocal fluorescence microscopy images of the proximal ascending aorta from 16-week-old *Lmna*^+/+^, *Lmna*^G609G/G609G^, and *Zmpste24*^−/–^ mice stained with antibodies against smooth muscle actin (Sm-actin, *green*), collagen type VIII (Col VIII, *red*), and CD31 (*magenta*). Nuclei were stained with Dapi (*white*). Scale bar, 50 µm. *Yellow* dotted lines mark the borders of the medial layer. *Red* arrowheads point to areas with reduced Sm-actin staining and reduced numbers of SMC nuclei. The *yellow* arrowhead points to collagen type VIII staining in the adventitia. Images of the entire tissue sections are shown in **Supplementary Figure 1A**. **B**. Confocal fluorescence microscopy images of the proximal ascending aorta from 13-week-old *Lmna*^+/+^ and *Lmna*^G609G/G609G^ mice and a 21-week-old *Zmpste24*^−/–^ mouse, all expressing a nuclear-targeted tdTomato transgene, stained with an antibody against farnesyl-prelamin A. The boxed regions are shown at higher magnification in the middle and far-right columns. The images show Dapi (*white*), elastic fibers (*green*), tdTomato (tdTom; *red*), and farnesyl-prelamin A (PreA; *magenta*). The *yellow* arrowhead (middle column) points to tdTomato outside of an SMC nucleus. Scale bar, 10 µm. Images of the entire tissue sections are shown in **Supplementary Figure 1B**. **C**. Diagram showing the location where SMC nuclei in the thoracic aorta were quantified (Proximal ascending, PA). **D–E**. Representative microscopy images of sections from the ascending thoracic aorta [stained with Dapi (*blue*)] from a 21-week-old *Zmpste24*^+/+^ mouse (**D**) and a *Zmpste24*^−/–^ mouse (**E**). Scale bar, 100 µm. **F**. Bar graph showing the numbers of nuclei (relative to area) in the proximal ascending thoracic aorta in 21-week-old *Zmpste24*^+/+^ (WT) and *Zmpste24*^−/–^ (ZmpKO) mice. Mean ± SEM (*n* = 5 mice/group). Student’s *t* test. *P* = 0.84. Images of the entire tissue sections used for quantification are shown in **Supplementary Figure 2**.

### Progerin and farnesyl-prelamin A have similar effects in cultured SMCs

We used doxycycline (Dox)-inducible constructs to create mouse SMC clones that expressed human versions of progerin (Prog-SMC), mature lamin A (PreA-SMC), and farnesyl-prelamin A (PreA-ZMPKO-SMCs). The levels of nuclear lamin expression in the SMC clones were adjusted to match the amounts of progerin in aortas of *Lmna*^G609G/+^ mice (**Fig. 2a**). The expression of farnesyl-prelamin A in PreA-ZMPKO-SMCs was confirmed by western blotting with monoclonal antibody (mAb) 3C8, which binds preferentially to the farnesylated form of prelamin A (37) (**Fig. 2a**).

**Fig. 2.**
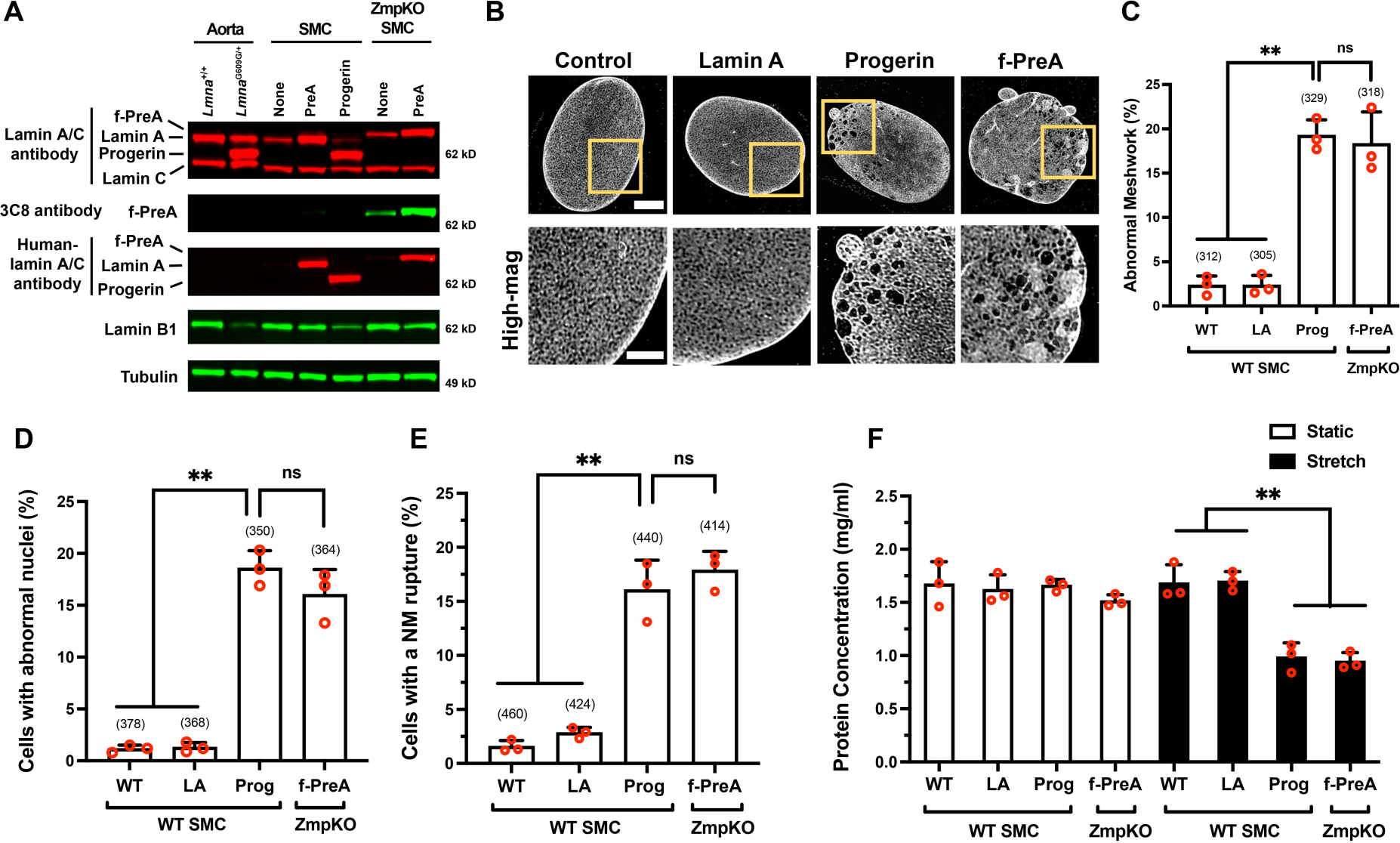
Progerin and farnesyl-prelamin A have similar effects in cultured SMCs. **A**. Western blot comparing the expression of nuclear lamins in the mouse aorta and in mouse SMCs expressing human lamin A, human progerin, and human farnesyl-prelamin A. Nuclear lamins were detected with antibodies that bind to mouse and human lamin A/C, farnesyl-prelamin A (clone 3C8), human lamin A/C, and lamin B1. Tubulin was measured as a loading control. **B**. Representative high-resolution confocal microscopy images showing the protein meshworks formed by human versions of lamin A, progerin, and farnesyl-prelamin A (f-PreA). Scale bar, 5 µm. The boxed regions are shown at higher magnification below. Scale bar, 2 µm. **C**. Nuclei with an abnormal meshwork in SMCs expressing human lamin A (LA), progerin (Prog), and farnesyl-prelamin A (f-PreA). Mean ± SEM (*n* = 3 experiments). ANOVA. **, *P* < 0.01. ns, not significant. **D**. Abnormal nuclear shape in SMCs expressing human versions of lamin A, progerin, and farnesyl-prelamin A. Mean ± SEM (*n* = 3 experiments). ANOVA. **, *P* < 0.01. ns, not significant. **E**. Nuclear membrane (NM) ruptures in SMCs expressing human versions of lamin A, progerin, and farnesyl-prelamin A. Mean ± SEM (*n* = 3 experiments). ANOVA. **, *P* < 0.01. ns, not significant. **F**. Cell death in SMCs expressing human versions of lamin A, progerin, or farnesyl-prelamin A. SMCs were cultured on PDMS membranes and exposed to static (open bars) or cyclical stretching conditions (closed bars) for 24 h. The fraction of cells remaining on the membranes were quantified by measuring protein concentration. Mean ± SEM (*n* = 3 experiments). ANOVA. **, *P* < 0.01. For the data reported in C–E, the number of nuclei or cells examined are shown in parentheses.

The nuclear lamin meshworks formed by progerin, farnesyl-prelamin A, and mature lamin A (in Prog-SMCs, PreA-ZMPKO-SMCs, and PreA-SMCs, respectively) were examined by high-resolution confocal fluorescence microscopy after staining with a human lamin A/C–specific antibody (31). The mature lamin A in PreA-SMCs formed a meshwork with small, uniform gaps and was morphologically indistinguishable from the mouse lamin A/C meshwork in wild-type SMCs (**Fig. 2b**). In contrast, progerin and farnesyl-prelamin A both formed an abnormal meshwork (with large and irregular-sized gaps) in SMCs (∼17% for both) (**Figs. 2b–c**). Aside from triggering similar morphological abnormalities in the nuclear lamin meshwork, progerin and farnesyl-prelamin A triggered very similar frequencies of misshapen nuclei and nuclear membrane (NM) ruptures (**Fig. 2d**). Misshapen nuclei were present in 18% of Prog-SMCs and in 16% of PreA-ZMPKO-SMCs but were present in less than 1% of PreA-SMCs (≥ 350 cells/group). Similarly, NM ruptures [detected by live-cell imaging (21)] were present in 16% of Prog-SMCs and 18% in PreA-ZMPKO-SMCs, but were present in less than 3% of PreA-SMCs (≥ 414 cells/group) (**Fig. 2e**). In cells subjected to cyclical stretching (15), progerin and farnesyl-prelamin A increased the frequency of cell death to a similar degree (**Fig. 2f**). Cell death was observed in 40% of Prog-SMCs, 38% of PreA-ZMPKO-SMCs, but in less than 1% of PreA-SMCs.

### Farnesyl-prelamin A does not accumulate with age in the aorta of *Zmpste24*^−/–^ mice

We hypothesized that the striking differences in aortic pathology in *Lmna*^G609G/G609G^ and *Zmpste24*^−/–^ mice (despite very similar toxicities of progerin and farnesyl-prelamin A in cultured SMCs) could be due to different amounts of progerin and farnesyl-prelamin A in aortas. To explore that possibility, we examined aortas of 5- and 21-week-old *Zmpste24*^−/–^ and *Lmna*^G609G/+^ mice (**Fig. 3a**). Consistent with earlier findings (21), the levels of progerin in the aortas of *Lmna*^G609G/+^ mice increased with age, whereas the levels of lamin B1 fell. Levels of progerin increased by 62% between 5 and 21 weeks of age, while lamin B1 levels fell by 57% (*n* = 4 mice/group) (**Fig. 3b**). The age-related increase in progerin could not be explained by higher transcript levels; progerin transcript levels in the aorta fell by 22% with age (**Fig. 3c**). *Lmnb1* transcripts in *Lmna*^G609G/+^ mice declined by 61% (**Fig. 3e**). In parallel, we examined aortas in age-matched *Zmpste24*^−/–^ mice (*n* = 8 mice/group). In young animals, levels of farnesyl-prelamin A in *Zmpste24*^−/–^ mice were equivalent to the progerin levels in *Lmna*^G609G/+^ mice. In contrast to the age-related increase in progerin levels in *Lmna*^G609G/+^ aortas, the levels of farnesyl-prelamin A in *Zmpste24*^−/–^ aortas decreased by 19% (**Fig. 3d**). Lamin B1 protein levels in *Zmpste24*^−/–^ aortas decreased by 53% (**Fig. 3d**). Prelamin A transcript levels in *Zmpste24*^−/–^ aortas fell by 15% at 21 weeks; *Lmnb1* transcript levels were reduced by 61% (**Fig. 3e**).

**Fig. 3.**
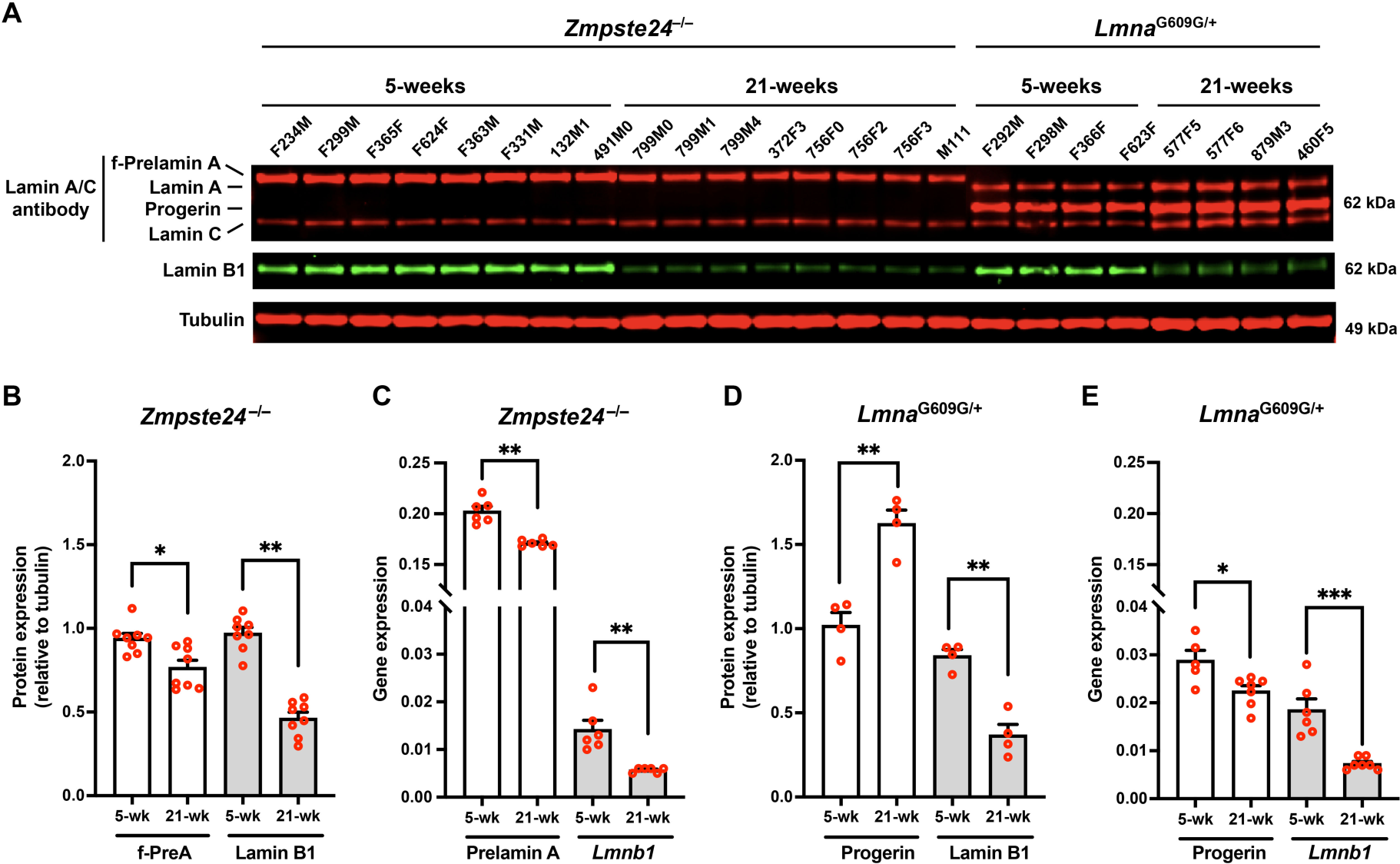
Farnesyl-prelamin A does not accumulate with age in the aorta of *Zmpste24*^−/–^ mice. **A**. Western blot comparing the expression of farnesyl-prelamin A, progerin, and lamin B1 in aortas from young and old *Zmpste24*^−/–^ and *Lmna*^G609G/+^ mice. Tubulin was measured as a loading control. The ages and mouse IDs are shown above each sample. **B**. Bar graph showing farnesyl-prelamin A (f-PreA) and lamin B1 expression (relative to tubulin) in aortas from young and old *Zmpste24*^−/–^ mice. Mean ± SEM (*n* = 8 mice/group). Student’s *t* test. *, *P* < 0.05. **, *P* < 0.01. **C.** Quantitative RT-PCR studies showing prelamin A and *Lmnb1* transcript levels in aortas from young and old *Zmpste24*^−/–^ mice. Mean ± SEM (*n* = 6 mice). Student’s *t* test, **, *P* < 0.01. **D**. Bar graph showing progerin and lamin B1 protein expression (relative to tubulin) in aortas from young and old *Lmna*^G609G/+^ mice. Mean ± SEM (*n* = 4 mice/group). Student’s *t* test. **, *P* < 0.01. **E**. Quantitative RT-PCR studies showing progerin and *Lmnb1* transcript levels in aortas from young and old *Lmna*^G609G/+^ mice. Mean ± SEM (*n* = 5–7 mice/group). Student’s *t* test. *, *P* < 0.05. ***, *P* < 0.001.

### Progerin causes accumulation of A-type nuclear lamins in the aorta and heart

The fact that progerin levels increased with age in *Lmna*^G609G/+^ aortas without a corresponding increase in progerin transcripts [(21) and **Figs. 3b–c**] raised the possibility that progerin itself alters the turnover of nuclear lamins. To explore that possibility, we isolated aortas from young (5 week) and old (14 week) *Lmna*^G609G/+^ mice and quantified aortic levels of lamin A, lamin C, and progerin (*i.e*., all of the A-type nuclear lamins in *Lmna*^G609G/+^ mice). The levels of all three A-type nuclear lamins increased with age (**Figs. 4a–b**). We also examined aortas in age-matched *Zmpste24*^−/–^ mice. The A-type nuclear lamins in *Zmpste24*^−/–^ mice (farnesyl-prelamin A and lamin C) did not increase with age (**Figs. 4a, 4c**). To test whether the expression of progerin was responsible for the age-related increase in A-type nuclear lamin levels in the aorta, we bred *Zmpste24*-deficient mice that also express progerin (*Zmpste24*^−/–^*Lmna*^G609G/+^). In contrast to the observations in *Zmpste24*^−/–^ mice, the levels of farnesyl-prelamin A and lamin C in *Zmpste24*^−/–^*Lmna*^G609G/+^ aortas increased with age (by 1.9- and 2-fold, respectively) (**Figs. 4a, 4d**). In these studies, aortas were examined at 14 weeks rather than 21 weeks of age because the disease phenotypes occurred earlier in *Zmpste24*^−/–^*Lmna*^G609G/+^ mice than in *Zmpste24*^−/–^ mice (**Supplementary Fig. 3a**). Quantitative PCR studies showed that prelamin A, progerin, and lamin C transcript levels in the aorta did not increase with age (**Supplementary Figs. 3b–d**). Thus, the age-related increase in A-type nuclear lamins in both *Lmna*^G609G/+^ and *Zmpste24*^−/–^*Lmna*^G609G/+^ aortas could not be explained by increased *Lmna* transcripts. Lamin B1 protein levels in aortas of *Lmna*^G609G/+^, *Zmpste24*^−/–^, and *Zmpste24*^−/–^ *Lmna*^G609G/+^ mice decreased with age (by 60%, 46%, and 79%, respectively; P < 0.01) (**Figs. 4e**).

**Fig. 4.**
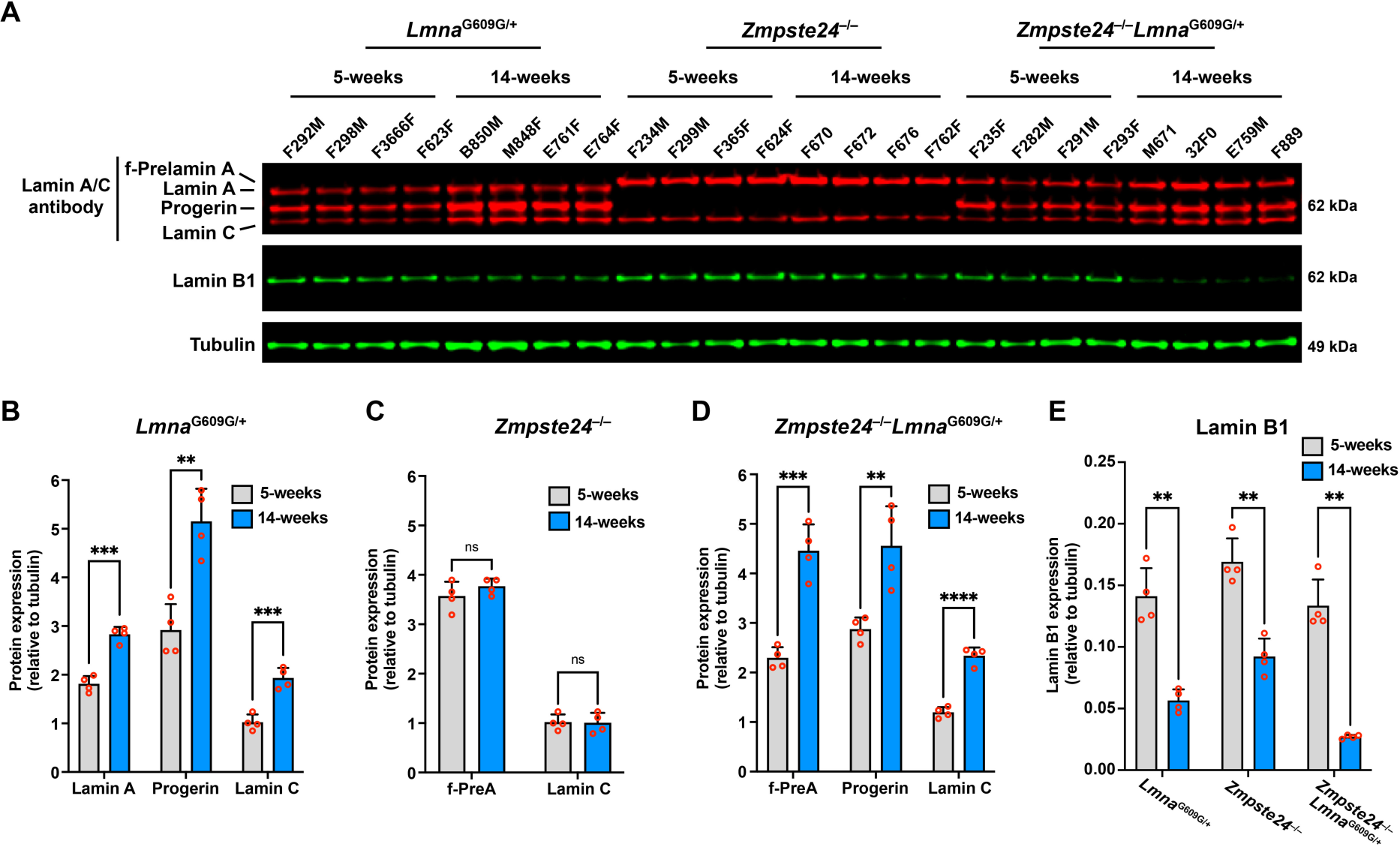
Progerin causes the accumulation of the A-type nuclear lamins in the aorta. **A**. Western blot comparing the expression of lamin A, lamin C, farnesyl-prelamin A, progerin, and lamin B1 in aortas from 5- and 14-week-old *Lmna*^G609G/+^, *Zmpste24*^−/–^, and *Zmpste24*^−/–^*Lmna*^G609G/+^ mice. Tubulin was measured as a loading control. The ages and mouse IDs are shown above each sample. **B**. Bar graph showing lamin A, progerin, and lamin C expression (relative to tubulin) in aortas from young and old *Lmna*^G609G/+^ mice. Mean ± SEM (*n* = 4 mice/group). Student’s *t* test. **, *P* < 0.01. ***, *P* < 0.001. **C**. Bar graph showing farnesyl-prelamin A (f-PreA) and lamin C expression (relative to tubulin) in aortas from young and old *Zmpste24*^−/–^ mice. Mean ± SEM (*n* = 4 mice/group). Student’s *t* test. ns, not significant. **D**. Bar graph showing f-PreA, progerin, and lamin C expression (relative to tubulin) in aortas from young and old *Zmpste24*^−/–^*Lmna*^G609G/+^ mice. Mean ± SEM (*n* = 4 mice/group). Student’s *t* test. **, *P* < 0.01. ***, *P* < 0.001. ****, *P* < 0.0001. **E**. Bar graph showing lamin B1 expression (relative to tubulin) in aortas from young and old *Lmna*^G609G/+^, *Zmpste24*^−/–^, and *Zmpste24*^−/–^*Lmna*^G609G/+^ mice. Mean ± SEM (*n* = 4 mice/group). Student’s *t* test. **, *P* < 0.01.

We also examined the impact of progerin on A-type nuclear lamins in the heart. Similar to the observations in aorta, amounts of A-type nuclear lamins in hearts of *Lmna*^G609G/+^ and *Zmpste24*^−/–^ *Lmna*^G609G/+^ mice increased with age, whereas the amounts remained stable in *Zmpste24*^−/–^ mice (**Supplementary Figs. 4a–d**). Lamin B1 levels in hearts of *Lmna*^G609G/+^ and *Zmpste24*^−/–^ *Lmna*^G609G/+^ mice decreased with age or remained the same (**Supplementary Fig. 4e**). Transcript levels for the A-type nuclear lamins in hearts of *Lmna*^G609G/+^ and *Zmpste24*^−/–^*Lmna*^G609G/+^ mice did not increase with age (**Supplementary Figs. 3e–g).**

### The loss of SMCs in aortas of *Zmpste24*-deficient mice can be induced by supraphysiologic levels of prelamin A production

We suspected that *Zmpste24*^−/–^ mice were protected from aortic SMC loss—despite the toxicity of farnesyl-prelamin A in cultured SMCs—was related to the absence of farnesyl-prelamin A accumulation with age. We reasoned that it might be possible to induce SMC loss in aortas of *Zmpste24*^−/–^ mice by increasing prelamin A production. To test this possibility, we bred *Zmpste24*^−/–^ mice harboring a prelamin A–only allele (*Lmna*^PLAO^), which channels all of the output from *Lmna* into prelamin A rather than into both lamin C and prelamin A (38). The levels of mature lamin A in the aorta of *Lmna*^PLAO/+^ mice (prelamin A in wild-type mice is processed to mature lamin A) were ∼20% higher than in wild-type mice; the levels in *Lmna*^PLAO/PLAO^ mice were increased by ∼40% (**Supplementary Figs. 5a–b**). To our surprise, the levels of farnesyl-prelamin A in aortas of *Zmpste24*^−/–^*Lmna*^PLAO/+^ mice did not change (**Supplementary Fig. 5c**). Further increasing prelamin A production by breeding *Zmpste24*^−/–^ *Lmna*^PLAO/PLAO^ mice was not a useful strategy because those mice die from progressive multisystem disease by ∼6–7 weeks of age (38).

To circumvent this roadblock, we bred *Sm22α*-*CreZmpste24*^fl/fl^*Lmna*^PLAO/PLAO^ mice to increase the levels of farnesyl-prelamin A selectively in SMCs. The expression of *Sm22α*-*Cre* in aortic SMCs was confirmed with a dual-fluorescence *Rosa*^nT-nG^ reporter allele where tdTomato is expressed in the nucleus of cells but switches to EGFP after *Cre*-recombination (39). That approach makes it possible to visualize both *Cre*-positive and *Cre*-negative cells in the same sample. As expected, the two-color reporter showed *Cre* expression in aortic SMCs of *Sm22α*-*Cre*-positive mice (**Supplementary Fig. 6a**). Consistent with that finding, farnesyl-prelamin A synthesis was detected in aortic SMCs by immunohistochemistry (**Supplementary Fig. 6b**). However, the two-color reporter also revealed that many SMCs in *Sm22α*-*Cre*-positive mice do not express *Cre* (*i.e.*, tdTomato-positive SMCs were still detected in the medial layer of the aorta). Thus, the expression of the *Sm22α*-*Cre* transgene in SMCs is variegated. Nonetheless, farnesyl-prelamin A levels were increased in *Sm22α*-*CreZmpste24*^fl/fl^*Lmna*^PLAO/PLAO^ mice. At 8 and 12 weeks of age, the levels of farnesyl-prelamin A in the aorta were ∼45% higher in *Sm22α*-*CreZmpste24*^fl/fl^*Lmna*^PLAO/PLAO^ mice than in *Zmpste24*^−/–^ mice. At 18 and 27 weeks of age, histopathology studies of *Sm22α*-*CreZmpste24*^fl/fl^*Lmna*^PLAO/PLAO^ mice revealed loss of aortic SMCs along with reduced smooth muscle actin staining (**Fig. 5c** and **Supplementary Figs. 7–8**). Thus, supraphysiologic levels of farnesyl-prelamin A in SMCs had triggered aortic pathology. Interestingly, the aortic pathology developed even though the levels of farnesyl-prelamin A in 20-week-old *Sm22α*-*CreZmpste24*^fl/fl^*Lmna*^PLAO/PLAO^ mice were lower than the levels at 12 weeks of age (**Figs. 5a–b**). The decreased farnesyl-prelamin A levels were accompanied by increased amounts of mature lamin A. The latter findings are almost certainly due to variegation in the expression of the *Sm22α*-*Cre* transgene in SMCs and a substantial competitive advantage of the subset of aortic SMCs that retained *Zmpste24* expression (see *Discussion*).

**Fig. 5.**
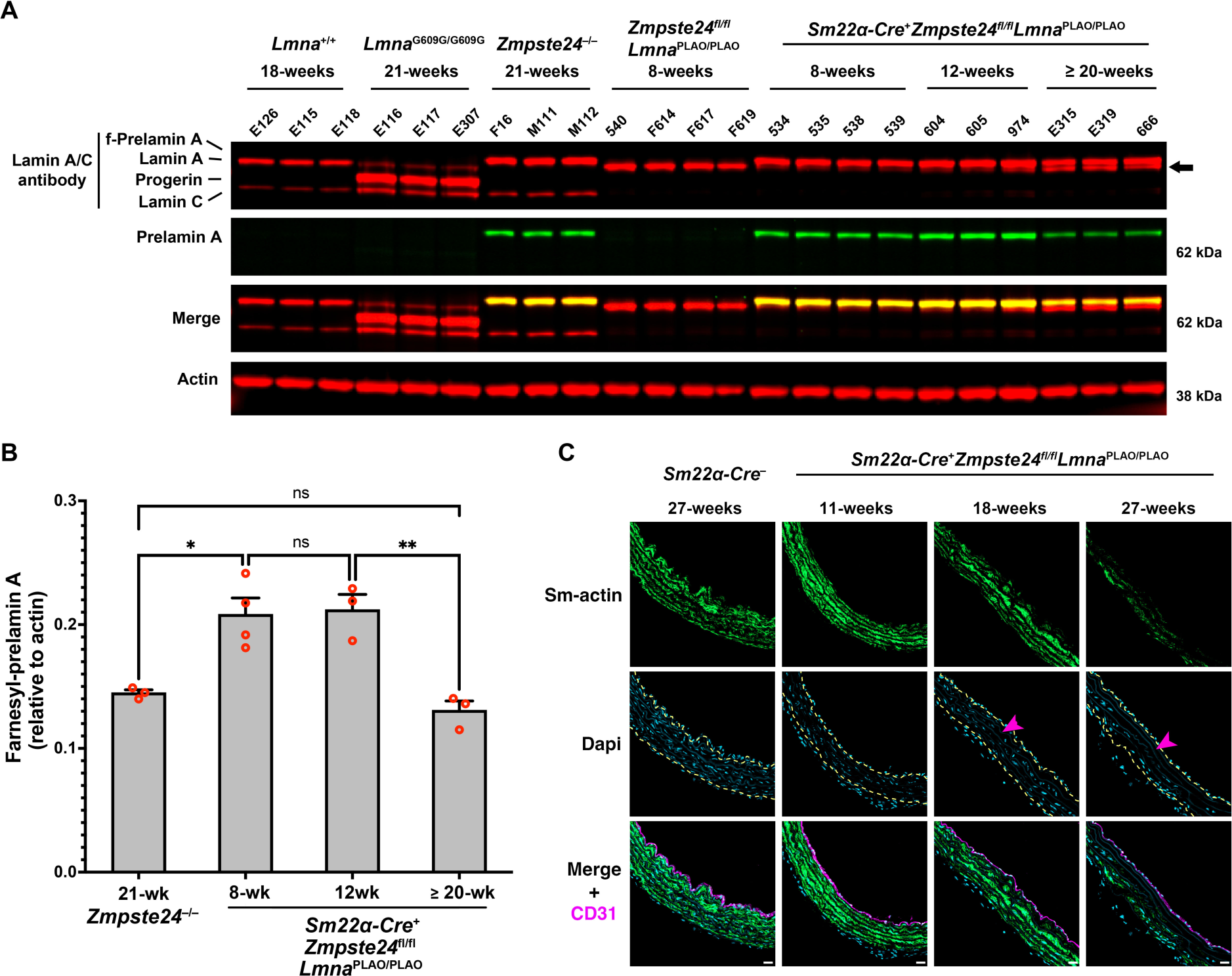
The loss of SMCs in aortas of *Zmpste24*-deficient mice can be induced by increasing prelamin A production. **A.** Western blot comparing the expression of lamin A, lamin C, farnesyl-prelamin A, progerin, and lamin B1 in aortas from wild-type, *Lmna*^G609G/G609G^, *Zmpste24*^−/–^, *Zmpste24*^fl/fl^*Lmna*^PLAO/PLAO^, and *Sm22α-CreZmpste24*^fl/fl^*Lmna*^PLAO/PLAO^ mice. Farnesyl-prelamin A was detected with monoclonal antibody 3C8. Actin was measured as a loading control. The ages and mouse IDs are shown above each sample. The arrow points to mature lamin A in older *Sm22α-CreZmpste24*^fl/fl^*Lmna*^PLAO/PLAO^ mice (detected with an anti-lamin A/C antibody). **B**. Bar graph comparing the expression of farnesyl-prelamin A (detected with antibody 3C8) relative to actin in 21-week-old *Zmpste24*^−/–^ mice and *Sm22α-CreZmpste24*^fl/fl^*Lmna*^PLAO/PLAO^ mice at 8, 12, and ≥ 20 weeks of age. ANOVA. \**P* < 0.05. \*\**P* < 0.01. ns, not significant. **C**. Confocal fluorescence microscopy images of the proximal ascending aorta from 11-, 18-, and 27-week-old *Sm22α-CreZmpste24*^fl/fl^*Lmna*^PLAO/PLAO^ mice stained with antibodies against smooth muscle actin (Sm-actin, *green*) and CD31 (*magenta*). As a control, images from a 27-week-old *Zmpste24*^fl/fl^*Lmna*^PLAO/PLAO^ mouse are shown. Nuclei were stained with Dapi (*blue*). Scale bar, 20 µm. *Yellow* dotted lines mark the borders of the medial layer. *Red* arrowhead points to an area with reduced Sm-actin staining and reduced numbers of SMC nuclei. Images of the entire sections are shown in **Supplementary Figure 7**. Images from additional 27-week-old *Sm22α-CreZmpste24*^fl/fl^*Lmna*^PLAO/PLAO^ mice are shown in **Supplementary Figure 8**.

### Reduced phosphorylation of serine-404 in progerin

Earlier studies established that phosphorylation of prelamin A at serine-404 by AKT triggers prelamin A degradation by lysosomal enzymes (40). Considering the differences in levels of progerin and farnesyl-prelamin A in older mice, we hypothesized that the age-related accumulation of progerin in *Lmna*^G609G/+^ mice might result from lower levels of serine-404 phosphorylation. To test this possibility, we compared the phosphorylation of serine-404 in human progerin and human farnesyl-prelamin A expressed in SMCs by western blotting (with an antibody against human lamin A/C phosphoserine-404). The specificity of the antibody was confirmed in cells expressing a S404A-human lamin A mutant and in *Lmna*^−/–^ SMCs (**Fig. 6a–b**). The phosphoserine lamin A antibody bound avidly to farnesyl-prelamin A in PreA-ZMPKO-SMCs, whereas the binding of the antibody to progerin in Prog-SMCs was very low (∼5% of the binding to farnesyl-prelamin A) (**Fig. 6a–b**). To determine if the reduced phosphorylation was due to the impact of progerin expression, we transiently expressed human prelamin A in ZMPKO-SMCs or in ZMPKO-SMCs that expressed progerin, and then examined the phosphorylation of farnesyl-prelamin A at serine-404. In the ZMPKO-SMCs expressing progerin, the phosphorylation of farnesyl-prelamin A was significantly reduced (**Figs. 6c–d**).

**Fig. 6.**
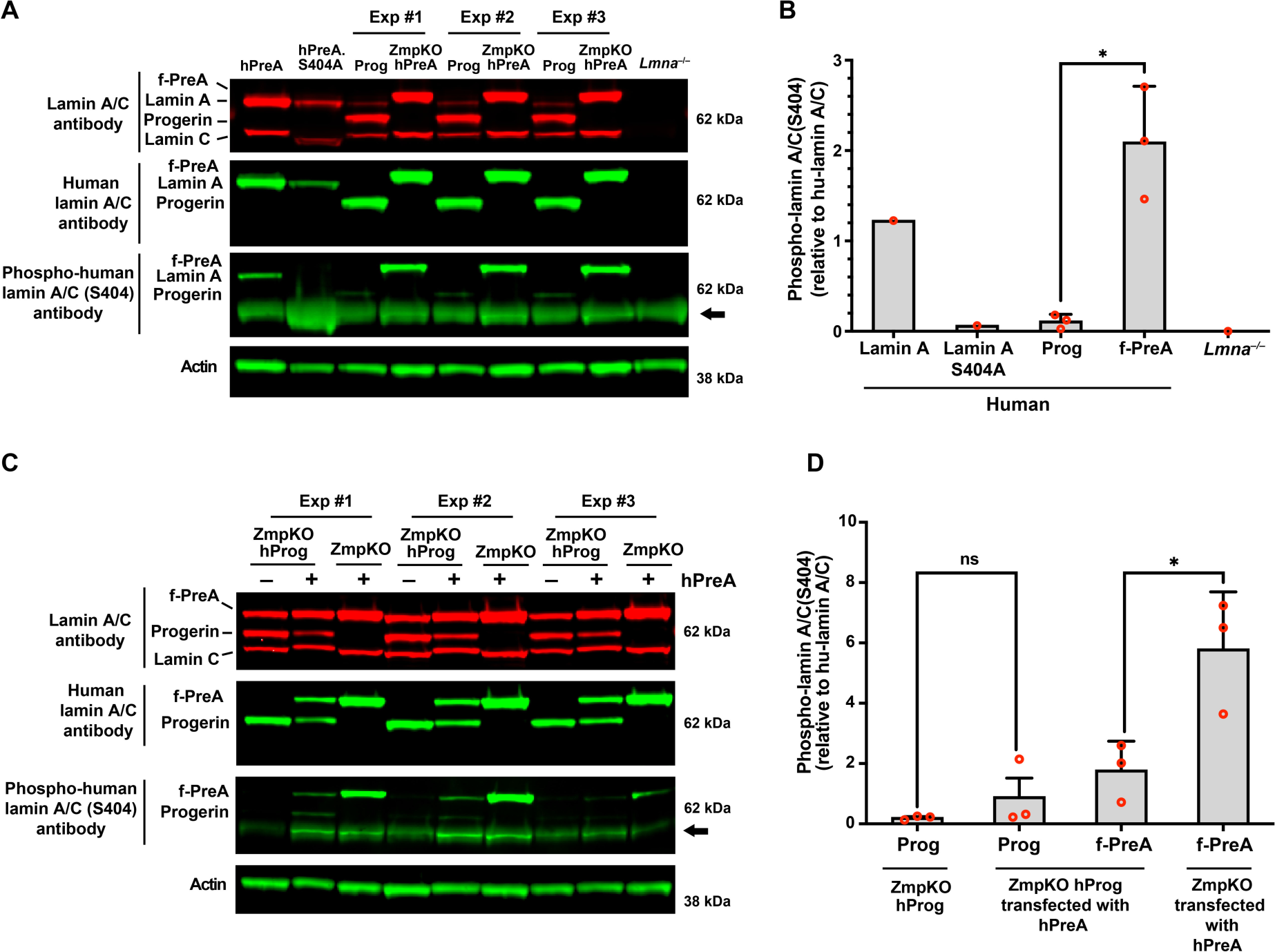
Reduced phosphorylation of serine-404 in progerin. **A**. Western blot comparing the levels of serine-404 phosphorylation in human versions of lamin A, farnesyl-prelamin A (f-PreA), and progerin (Prog) expressed in mouse SMCs. Serine-404 phosphorylation was detected with an antibody against human lamin A/C phosphoserine-404. The specificity of the antibody was evaluated using extracts from SMCs expressing an S404A human lamin A mutant and *Lmna*^−/–^ SMCs. The levels of expression of the human nuclear lamin proteins were measured with an anti-human lamin A/C antibody. Actin was measured as a loading control. The arrow points to a nonspecific band produced with the human lamin A/C phosphoserine-404 antibody. **B**. The bar graph compares the levels of serine-404 phosphorylation in the human versions of lamin A, lamin A-S404A, Prog, and f-PreA. Serine-404 phosphorylation was normalized to the level of human lamin protein expression. Mean ± SEM (*n* = 3). Student’s *t* test. *, P < 0.05. **C**. Western blot comparing the effects of progerin expression on the phosphorylation of serine-404 in human farnesyl-prelamin A. Human farnesyl-prelamin A synthesis was induced by transient transfection of human-prelamin A (hPreA) in *Zmpste24*^−/–^ SMCs or *Zmpste24*^−/–^ SMCs expressing progerin (ZmpKO hProg). Serine-404 phosphorylation in human farnesyl-prelamin A and progerin was detected as described in panel A. The arrow points to a nonspecific band produced with the human lamin A/C phosphoserine-404 antibody. **D**. The bar graph compares the effects of progerin expression on the levels of serine-404 phosphorylation in farnesyl-prelamin A. The levels of serine-404 phosphorylation were measured as described in panel B. Mean ± SEM (*n* = 3). ANOVA. *, P < 0.05. ns, not significant.

To determine if AKT activity is reduced in the aorta of *Lmna*^G609G/+^ mice, we compared the phosphorylation of AKT at serine-473 (41) in young and old wild-type (*Lmna*^+/+^), *Lmna*^G609G/+^, and *Zmpste24*^−/–^ mice. Relative to total amounts of AKT, AKT phosphorylation at serine-473 was significantly lower in *Lmna*^G609G/+^ mice than in age-matched *Lmna*^+/+^ and *Zmpste24*^−/–^ mice (**Fig. 7**). Consistent with reduced AKT activity in cells expressing progerin, AKT phosphorylation was also lower in 14-week-old *Zmpste24*^−/–^*Lmna*^G609G/+^ mice than in age-matched *Zmpste24*^−/–^ mice (reduced by > 90%; ANOVA. P < 0.01).

**Fig. 7.**
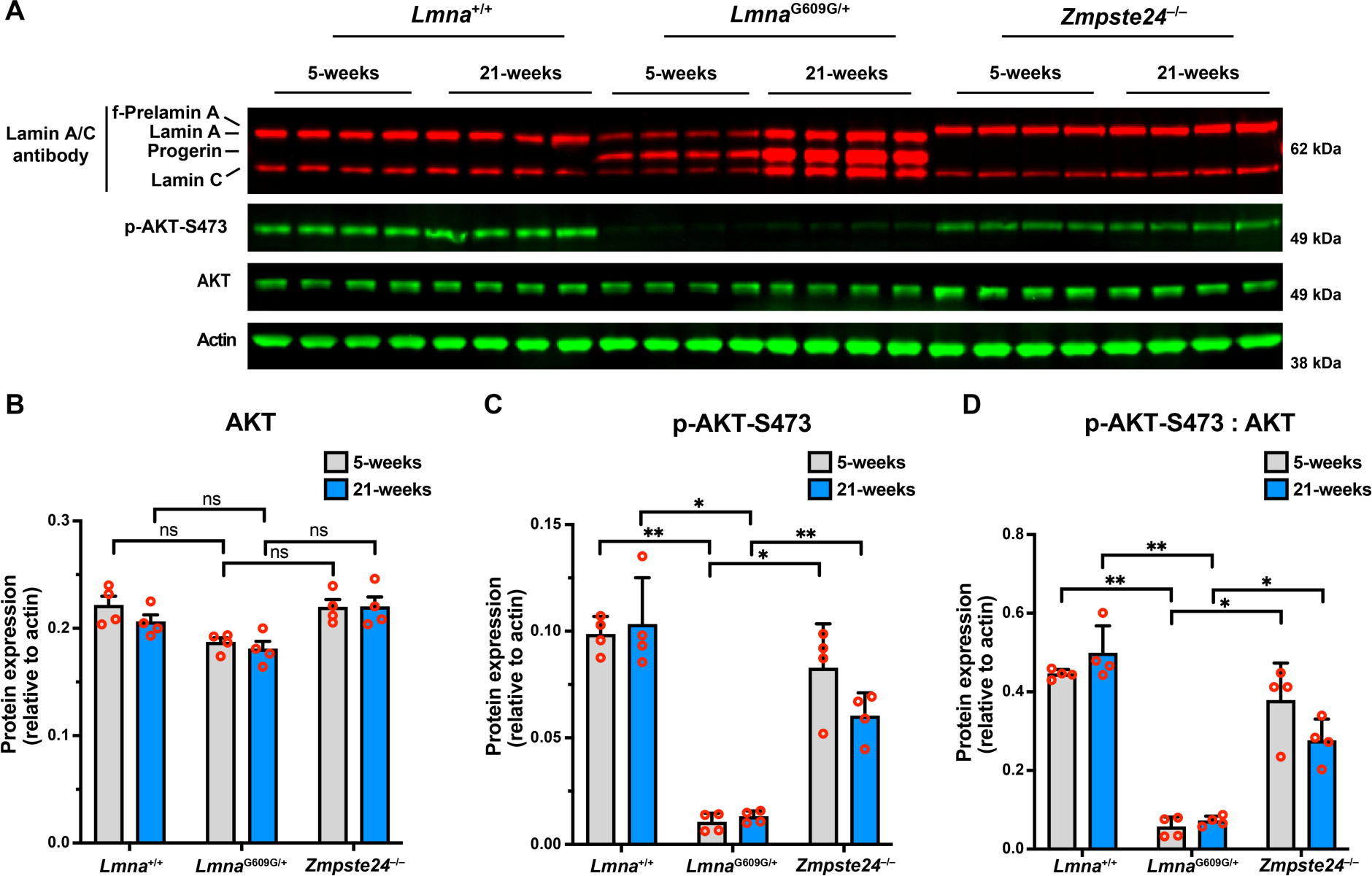
AKT activity is reduced in the aorta of *Lmna*^G609G/+^ mice. **A**. Western blot comparing the levels of phosphorylated AKT at serine-473 (p-AKT-S473) in aortas from young and old *Lmna*^+/+^, *Lmna*^G609G/+^, and *Zmpste24*^−/–^ mice. Actin was measured as a loading control. The ages and mouse IDs are shown above each sample. **B**. Bar graph shows the levels of total AKT (relative to actin) in young and old *Lmna*^+/+^, *Lmna*^G609G/+^, and *Zmpste24*^−/–^ mice. Mean ± SEM (*n* = 4 mice/group). ANOVA. ns, not significant. **C**. Bar graph shows the levels of p-AKT-S473 (relative to actin) in young and old *Lmna*^+/+^, *Lmna*^G609G/+^, and *Zmpste24*^−/–^ mice. Mean ± SEM (*n* = 4 mice/group). ANOVA. *, P < 0.05. **, P < 0.01. **D**. Bar graph shows the levels of p-AKT-S473 (relative to total AKT) in young and old *Lmna*^+/+^, *Lmna*^G609G/+^, and *Zmpste24*^−/–^ mice. Mean ± SEM (*n* = 4 mice/group). ANOVA. *, P < 0.05. **, P < 0.01.

## Discussion

The fact that progerin, an internally truncated and farnesylated prelamin A, triggers SMC loss in large arteries has been well documented (14, 16, 17). The causal relationship between progerin expression and SMC loss is substantiated by the observation that the extent of SMC loss is correlated with the amount of progerin expression (15, 25) and by the observation that progerin-triggered SMC loss in mice can be reversed by extinguishing progerin synthesis (42). Despite this progress, the reason that progerin triggers SMC loss in arteries but spares cells in other tissues has remained unclear. Also unclear is why SMC loss occurs in mouse models that express progerin but has not been observed in *Zmpste24*-deficient mice or *Lmna*^L648R/L648R^ mice (15, 36), which express a mutant full-length farnesyl-prelamin A that is resistant to ZMPSTE24 cleavage. To gain insights into that difference, we compared the effects of progerin and full-length farnesyl-prelamin A in both cultured cells and genetically modified mice. We found that progerin and farnesyl-prelamin A, when expressed in identical amounts in mouse SMCs, induce virtually identical levels of toxicity (as judged by the morphology of nuclear lamin meshworks and by the frequencies of misshapen cell nuclei, nuclear membrane ruptures, and cell death during mechanical stretching). In contrast to the cell culture findings, the impacts of progerin and farnesyl-prelamin A expression in genetically modified mice are distinct. In young mice, the levels of progerin in aortas of “HGPS mice” (*Lmna*^G609G^) and the levels of farnesyl-prelamin A in *Zmpste24*^−/–^ mice are similar. In older mice, however, progerin expression in *Lmna*^G609G^ mice resulted in an age-related accumulation of progerin (along with lamin A and lamin C), whereas in *Zmpste24*^−/–^ mice, the levels of farnesyl-prelamin A (and lamin C) did not accumulate with age. The accumulation of progerin could not be ascribed to changes in transcript levels, which either remained stable or declined with age, implying that progerin retards the turnover of A-type nuclear lamins. Consistent with that idea, AKT activity, which phosphorylates A-type nuclear lamins at serine-404 (43) and targets prelamin A for degradation (40), is reduced in the aortas of *Lmna*^G609G/+^ mice. These studies identify a distinct property of progerin that underlies the high levels of progerin in the aorta and the progerin-induced SMC loss in HGPS.

Our earlier studies identified an age-related accumulation of progerin in aortas of *Lmna*^G609G/+^ mice and raised the possibility of reduced progerin turnover (21). Building on that observation, we have, in the current study, shown that progerin levels in the aorta (and heart) not only accumulated with age but that this accumulation was accompanied by progerin-induced accumulation of the other A-type nuclear lamins. In contrast, full-length farnesyl-prelamin A in *Zmpste24*^−/–^ mice did not affect the levels of A-type nuclear lamins. In a recent report (44), Hasper and colleagues examined “lifetimes” of nuclear lamins in a knock-in mouse model of HGPS. They found that the turnover of progerin in the aorta was reduced in the HGPS mice, consistent with our results. However, they did not detect an accumulation of progerin. In those studies, they measured the abundance of all A-type nuclear lamins (lamin A, lamin C, progerin) as a group in both young (*n* = 3) and old (*n* = 2) HGPS mice. While aortic levels of the A-type lamins tended to be higher in HGPS mice than in wild-type mice, this difference was not judged to be significant “due to variability across samples.” Higher levels of the A-type nuclear lamins were observed in the heart of HGPS mice, but since the A-type nuclear lamins were measured as a group, they could not determine whether the higher levels were due to an accumulation of progerin alone, or to lamin A and lamin C, or to all three nuclear lamins.

We suspected that the absence of vascular pathology in *Zmpste24*^−/–^ mice was due to the fact that the levels of farnesyl-prelamin A in *Zmpste24*^−/–^ aortas were lower than the levels of progerin in *Lmna*^G609G/G609G^ aortas. To test whether increasing the levels of farnesyl-prelamin A expression were capable of eliciting aortic pathology, we bred *Zmpste24*^−/–^*Lmna*^PLAO/+^ mice. Because those mice die earlier than *Zmpste24*^−/–^ mice (38), we suspected that they might have significantly higher levels of farnesyl-prelamin A in the aorta, but this was not the case. The levels of farnesyl-prelamin A did not change. This was surprising given that the *Lmna*^PLAO^ allele increased the levels of lamin A in the aortas of wild-type mice (*i.e.*, 20% with one *Lmna*^PLAO^ allele and 40% with two *Lmna*^PLAO^ alleles). Breeding *Zmpste24*^−/–^ mice with two *Lmna*^PLAO^ alleles was not an option because those mice die by ∼6-weeks of age (38). We therefore produced SMC-specific *Zmpste24* knockout mice harboring two *Lmna*^PLAO^ alleles (*Sm22α*-*CreZmpste24*^fl/fl^*Lmna*^PLAO/PLAO^). At 8 and 12 weeks of age, the aortic levels of farnesyl-prelamin A in those mice were ∼45% higher than in *Zmpste24*^−/–^ mice. At 18 weeks of age, we observed substantial loss of aortic SMCs in *Sm22α*-*CreZmpste24*^fl/fl^*Lmna*^PLAO/PLAO^ mice, demonstrating that supraphysiologic levels of farnesyl-prelamin A are toxic to aortic SMCs *in vivo*. Unexpectedly, the levels of farnesyl-prelamin A did not remain stable. Instead, there was a decrease in farnesyl-prelamin A levels, accompanied by increased amounts of mature lamin A, in older *Sm22α*-*CreZmpste24*^fl/fl^*Lmna*^PLAO/PLAO^ mice. We suspect that those findings were secondary to the variegated expression of the *Sm22α*-*Cre* transgene. Because of the incomplete expression of the transgene in aortic SMCs, the aortas of *Sm22α*-*CreZmpste24*^fl/fl^*Lmna*^PLAO/PLAO^ mice contain a mixture of wild-type and *Zmpste24*-deficient SMCs. We believe that the supraphysiologic levels of farnesyl-prelamin A in the *Zmpste24*-deficient SMCs led to the death of those cells, resulting in increased numbers of SMCs expressing mature lamin A. Consistent with this interpretation, western blot studies revealed an age-related fall in farnesyl-prelamin A levels accompanied by increased levels of mature lamin A.

Initially, we suspected that the differences in vascular pathology in *Lmna*^G609G^ and *Zmpste24*^−/–^ mice could be related to different amounts of lamin B1 in SMCs. In earlier studies (15, 21), we observed that lamin B1 levels in the mouse aorta are very low. In wild-type mice, the low levels of lamin B1 remained stable with advancing age, while in *Lmna*^G609G^ mice, the lamin B1 levels fell with increasing age. In cultured cells, we observed that increased expression of lamin B1 reduces some of the toxic effects of progerin (*e.g.*, abnormal meshwork, nuclear membrane ruptures, stress-induced cell death), whereas reducing lamin B1 expression has the opposite effect (15, 21, 31). Given these observations, we suspected that we might find significantly higher aortic levels of lamin B1 in *Zmpste24*^−/–^ mice than in *Lmna*^G609G^ mice, but this was not the case. The aortic levels of lamin B1 were low in *Zmpste24*^−/–^ mice, and the levels in *Zmpste24*^−/–^and *Lmna*^G609G/+^ mice declined with age to a similar degree. Instead, we found that the greater susceptibility of *Lmna*^G609G^ mice to vascular pathology was due to a greater age-related accumulation of progerin. Although the levels of the other A type nuclear lamins also increased with age in *Lmna*^G609G^ mice (which might be expected to mitigate disease in the aorta), it is likely that the dominant-negative effects of progerin limit any beneficial effects provided by the increased levels of lamin A and lamin C. Progerin levels also increased in the heart with age, but in contrast to the aorta, we could not detect an extensive loss of cardiomyocytes in the heart. The key difference between the heart and the aorta is that the levels of *Lmna* expression are far lower in the heart. Thus, even though progerin accumulates with age in the heart of *Lmna*^G609G/G609G^ mice, the levels of progerin in the heart never approach the extremely high levels observed in the aorta. Quantitative western blots have demonstrated that the progerin-to-lamin B1 ratio is ∼10-fold higher in the aorta than in the heart (15).

The phosphorylation/dephosphorylation of nuclear lamins (45) has long been recognized to be crucial for the disassembly and assembly of the nuclear lamina during mitosis [reviewed in (46)], but nuclear lamin phosphorylation also affects the interaction of nuclear lamins with other proteins, the subcellular localization of nuclear lamins, and nuclear lamin levels (47, 48). The phosphorylation of serine-404 in prelamin A by AKT affects the localization of prelamin A and triggers its degradation in lysosomes (40, 43). Progerin is also phosphorylated at serine-404 (48), but it has not been clear whether that phosphorylation regulates progerin levels. Given that AKT activity is perturbed in both cell culture and mouse models of HGPS (41, 49), we hypothesized that the age-related accumulation of progerin in the aorta could be due to reduced phosphorylation by AKT. In cultured cells, serine-404 was strongly phosphorylated in lamin A and farnesyl-prelamin A, but weakly phosphorylated in progerin. The reduced phosphorylation could theoretically be due to differences in protein structure (as a consequence of the missing 50-amino acid segment) (50), thereby reducing AKT-dependent phosphorylation, but the fact that progerin expression also reduced the phosphorylation of farnesyl-prelamin A suggested that progerin might affect AKT activity. We were encouraged by these findings and hoped to investigate the phosphorylation of progerin in the aorta of mice, but antibodies against mouse lamin A/C phosphoserine-404 were not available. We did, however, examine AKT activity as judged by AKT phosphorylation at serine-473 (41, 49). AKT activity was lower in aortas of *Lmna*^G609G/+^ mice than in aortas of wild-type or *Zmpste24*^−/–^ mice. Consistent with the ability of progerin to reduce AKT activity, AKT phosphorylation in aortas of *Zmpste24*^−/–^*Lmna*^G609G/+^ mice was lower than in aortas of *Zmpste24*^−/–^ mice.

Our studies suggest a model for the loss of aortic SMCs in HGPS (**Fig. 8**). In SMCs of the mouse aorta, *Lmna* expression is high and *Lmnb1* expression is low. In mouse models of HGPS, this expression pattern translates into high levels of progerin and low levels of lamin B1 in aortic extracts (15). This high progerin:lamin B1 ratio results in gaps and irregularities in the nuclear lamina meshwork, reduced structural integrity of the nuclear lamina, and increased susceptibility to mechanical stress (15, 21, 31). As the mice age, the progerin:lamin B1 ratio in the aorta is even higher (21), a consequence of both progerin accumulation and declining levels of lamin B1. That combination leads to frequent nuclear membrane ruptures, DNA damage, and ultimately to the loss of aortic SMCs.

**Fig. 8.**
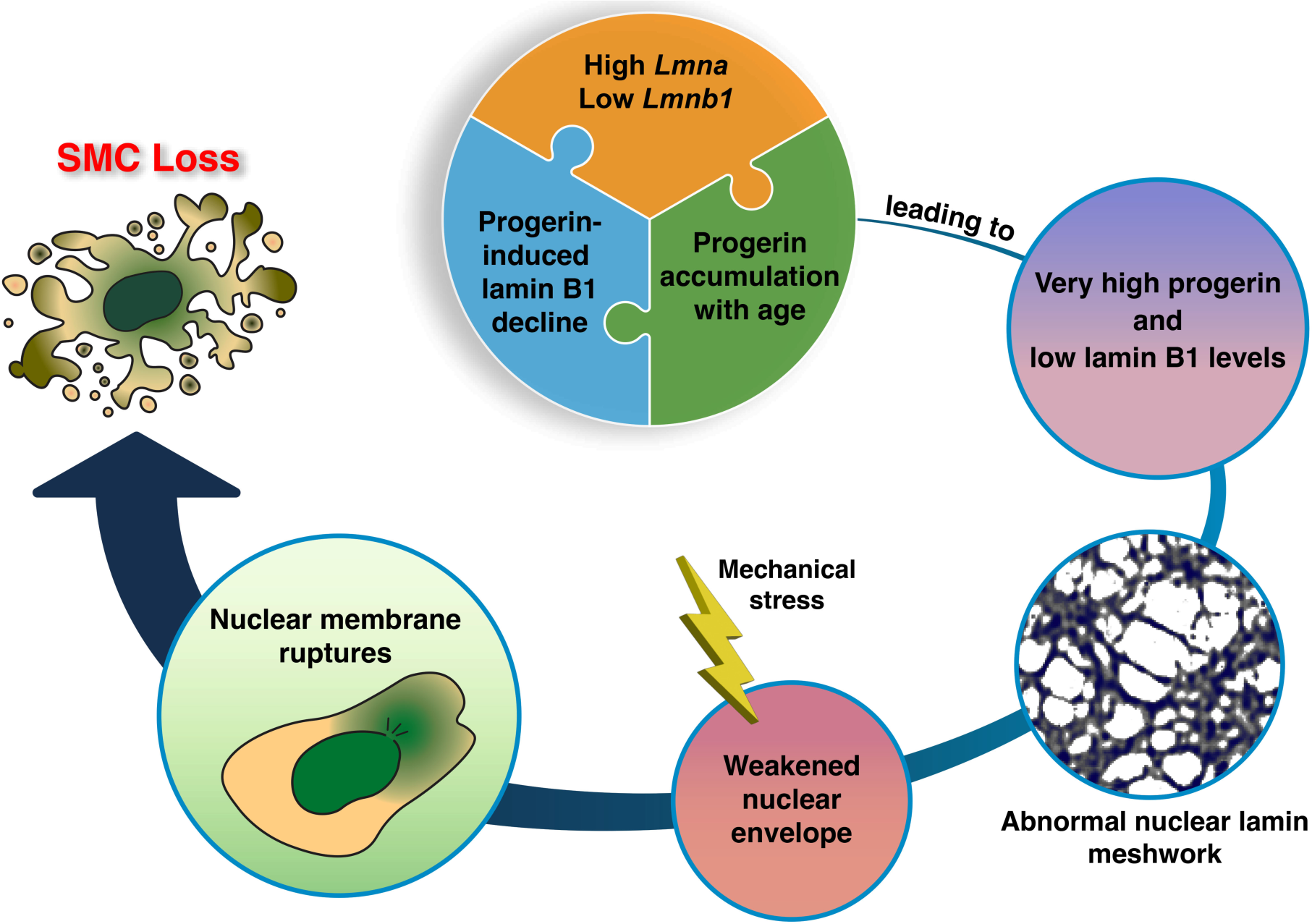
SMC loss in a mouse model of HGPS. A high *Lmna* and low *Lmnb1* expression profile in the mouse aorta results in a high lamin A:lamin B1 protein profile—ranking at the very top compared to 10 other mouse tissues (*e.g.*, ∼10-fold higher than in the kidney) (15). In the setting of HGPS, the high progerin:lamin B1 ratio increases with age and reaches extremely high levels (*e.g.*, ∼30-fold higher than in the kidney) (15). This increase is due to two factors. First, progerin, the toxic molecule in HGPS, increases the turnover of lamin B1, thereby reducing lamin B1 levels in the aorta (21). Lamin B1 acts in vitro to reduce the toxicity of progerin (21, 31). Second, progerin causes the accumulation of progerin itself as well as the other A-type nuclear lamins (shown in the current studies). We propose that the very high progerin:lamin B1 profile in aortic SMCs underlies the morphologically abnormal nuclear lamin meshwork and the reduced integrity of the nuclear envelope (31). In the setting of HGPS and mechanical stress (*e.g*., pulsatile flow in the aorta), the structural abnormalities in the nuclear lamin meshwork lead to nuclear membrane ruptures, DNA damage, and ultimately to loss of SMCs in the large arteries (15, 21).

## Methods

### Mice

*Zmpste24*^−/–^ (51), *Lmna*^PLAO^ (38), *Lmna*^G609G^ (17), and *Zmpste24*^fl/fl^ (37) mouse strains have been described previously. The *Sm22α-Cre* (stock no. 017491) and ROSA^nT-nG^ (stock no. 023035) mouse strains were purchased from The Jackson Laboratory (Bar Harbor, ME). *Zmpste24*^−/–^ *Lmna*^G609G/+^ (and littermate *Zmpste24*^−/–^ mice) were produced by intercrossing *Zmpste24*^+/–^ *Lmna*^G609G/+^ mice. All mice were housed in a specific pathogen–free barrier facility with a 12-h light/dark cycle. The mice were provided pelleted mouse chow (NIH31) and water *ad libitum*. Both male and female mice were used for experimental studies.

### Immunohistochemical analysis of aortic tissue

Mice were perfused in situ with PBS followed by fixative solution (3% paraformaldehyde in PBS). The thoracic aorta was dissected free and incubated in fixative solution at 4°C for 1–2 h. Aortic rings (∼2-mm) from the proximal ascending aorta, proximal descending, and mid-descending aorta were embedded in OCT, and frozen sections (4–6-µm) collected onto glass slides. Tissue sections were incubated as described previously (15, 21) with antibodies and concentrations listed in **Supplementary Table 1**. Nuclei were stained with Dapi. The stained sections were coded, and images were captured on a Zeiss LSM800 confocal microscope with a Plan-Apochromat 20×/0.8 NA objective. The coded images (tiff format) were imported into ImageJ, and the numbers of nuclei in the media were counted by trained observers and expressed relative to media area.

### *Zmpste24*-deficient SMCs

Guide RNAs targeting exon 6 of *Zmpste24* (5′-GGTAAGGCTACCTGGAGGTG-3′) and (5′-ACAACATACACCTTAGTCAA-3′) were designed with Synthego’s gRNA design tool. Double-stranded gRNAs were subcloned into pX458-GFP CRISPR/Cas9 vector linearized with *Bbs*I. A Nucleofector II apparatus (Lonza) and the Cell line T Nucleofector kit (Lonza) were used to electroporate 2 µg of pX458 vectors containing the gRNAs into 2 × 10^6^ SMCs. After 48 h, transfected SMCs were cell sorted for the top 10% of GFP intensity by flow cytometry. Individual clones (20–30) were isolated by limiting dilution. Genomic DNA was extracted from SMC clones with the DNeasy kit (Qiagen) and subjected to PCR analysis. PCR primers outflanking the gRNA cut sites (5′-ATTGCCTGTGTCTGCCCTTCTGCT-3′ and 5′-GAACACTGGTTTTGTTTTGCAGCC-3′) were used to amplify the gene fragment from the genomic DNA. Sequencing primer (5′-TCCACCAGCATGAACAAGGGTGTGT-3′) was used to verify the sequence deletion between the two guide RNA cut sites. Clonal cell lines were further tested by qPCR and western blotting to confirm the absence of ZMPSTE24 activity.

### *Lmna*-deficient SMC

*Lmna-*deficient SMCs were generated in a similar fashion as *Zmpste24*-deficient SMCs. Guide RNAs targeting the 5′ and 3′ UTRs of *Lmna* (5′-GGATTGGCCGCTTCTGTGCG-3′) and (5′-CCAATCGCCGCACCTCTAGA-3′) were designed. Individual clones were isolated, and genomic DNA was extracted for sequencing. Deletion of the entire *Lmna* gene was confirmed by sequencing, qPCR, immunofluorescence, and western blotting.

### Measurement of nuclear membrane (NM) ruptures in live SMCs

SMCs stably expressing nls-GFP were seeded into 2-well chamber slides with #1.5 glass coverslip bottom (ThermoFisher Scientific) and cultured in complete media. To induce cell-cycle arrest, mitomycin C (1 µg/ml) was added to the media. The inhibition of mitosis greatly facilitated quantification of NM ruptures. Doxycycline was added to induce nuclear lamin expression for 24 h before examining the cells by confocal microscopy. The chamber slides were placed on a Zeiss LSM800 confocal laser-scanning microscope with a CO₂ and temperature-controlled stage, operated with Zen Blue 2.3 software (Zeiss). Cells were imaged for 48 h at 37°C with 5% CO₂ using a Plan-Apochromat 20×/0.8 NA objective. Images from randomly selected fields were captured every 1–2 min. The ratio of cells with a NM rupture, identified by the presence of GFP in the cytoplasm, was calculated by dividing the number of cells with a rupture by the total number of cells in the field.

### High-resolution confocal fluorescence microscopy

A total of 50,000 cells were seeded in a chambered slide with a #1.5H (170 µm ± 5 µm) glass bottom (ibidi USA). After 48 h, the cells, with or without doxycycline treatment, were fixed using 4% paraformaldehyde in PBS for 10 min at room temperature, followed by permeabilization with 0.3% Triton X-100 in PBS for 10 min. Immunofluorescence microscopy was performed as previously described (31). The antibodies and their concentrations are detailed in **Supplementary Table 1**. Airyscan images were captured using a Zeiss LSM980 equipped with Airyscan2 in super-resolution (SR) imaging mode, with a scan speed of 5 and with a Plan-Apochromat 63×/1.4 NA oil-immersion objective. The excitation wavelengths and filter settings for each dye were chosen based on the integrated dye presets in ZEN Blue 2.3 software, and consistent settings were maintained throughout the imaging session. *Z*-stacks were acquired at an optimal section thickness of 0.14 µm, starting from near the glass bottom to the top of the nucleus. Images collected in SR mode were further enhanced with Airyscan Joint Deconvolution (Zeiss) with the following parameters: sample structure set to standard, maximum iterations at 10, and a quality threshold of 0.00. SR confocal fluorescence images were processed with ZEN Blue 2.3 software to create maximum intensity projection images from the equatorial plane to the top of the nucleus (furthest from the glass bottom).

### Quantification of lamina meshwork gap

*Z*-axis images from the equator to the top of a nucleus were compiled and converted to 8-bit grayscale in ImageJ (NIH). Minor adjustments to brightness and contrast were made to optimize visualization of the nuclear lamina meshwork. The meshwork was outlined with the “Overlay” tool with a paintbrush set to 1-pixel width. A threshold was applied to define the meshwork, and the “Analyze Particles” function was used to quantify the areas of the gaps within the meshwork. A global scale was set using the original image’s scale bar. Meshwork gaps adjacent to the image borders were excluded from the analysis. The distribution of lamina meshwork gap areas was plotted for each nucleus. Nuclei exhibiting gap sizes greater than five times the average gap area were classified as abnormal.

### Measurement of cell death in stretched SMCs

Wild-type SMCs (1 × 10^5^) expressing human prelamin A, SMCs expressing human progerin, or ZMPSTE24-deficient SMCs expressing human prelamin A were seeded onto polydimethylsiloxane (PDMS) membranes and cultured for 48 h. The membranes were clamped in a custom-built biaxial cell stretching device (15) and stretched 3-mm at 0.5 Hz for 24 h. To measure protein concentration of surviving cells, membranes were rinsed with PBS and digested with 0.1 N NaOH. Protein content was measured with the DC protein assay kit (Bio-Rad).

### Statistical analysis

Statistical analyses were performed with Microsoft Excel for Mac 2021 and GraphPad Prism software. Experimental groups were analyzed by unpaired 2-tailed Student’s *t* test, or one-way and two-way ANOVA with Tukey’s multiple comparisons test. Statistical differences were considered significant when the P value was < 0.05. Red circles in bar graphs show the average values of independent experiments or values for individual animals.

### Study approval

All animal studies were approved by UCLA’s Animal Research Committee.

## Data availability

All data are available in the main text or the Supplementary Materials.

## Author contributions

PHK, SGY, and LGF designed the research studies. PHK, JRK, PJH, HJ, YT, AP, JS, RGY, LGF performed the experiments. PHK, SGY, and LGF wrote the first draft of the manuscript. PHK, JRK, SGY, and LGF edited the manuscript.

## Acknowledgments

We thank Dino Di Carlo (UCLA) for the use of the plasma cleaner. This work was supported by the National Institutes of Health grants HL171737 and HL139725.

## Supplementary Materials

### Materials and Methods

#### Culture of aortic smooth muscle cells

Immortalized mouse aortic smooth muscle cells (SMCs) were purchased from ATCC (#CRL-2797) and cultured in DMEM (Invitrogen) supplemented with 10% (v/v) fetal bovine serum (HyClone), 1× nonessential amino acids, 2 mM glutamine, 1 mM sodium pyruvate, and 0.2 mg/ml G418 at 37°C with 5% CO_2_.

#### Western blotting

Urea-soluble protein extracts from tissues and SMCs were prepared as described previously (*17, 21*). The extracts were mixed with LDS sample buffer (Invitrogen) and heated at 70°C for 10 min. The samples were size-fractionated on 4–12% gradient polyacrylamide Bis-Tris gels (Invitrogen) and transferred to nitrocellulose membranes. The membranes were blocked with Odyssey Blocking solution (LI-COR Bioscience, Lincoln, NE) for 1 hour at RT and incubated with primary antibodies at 4°C overnight. After washing the membranes with PBS containing 0.2% Tween-20 (3 times for 10 min each), they were incubated with infrared (IR) dye– labeled secondary antibodies at RT for 1 hour. Membranes were washed with 0.2% PBS-T (3 times for 10 min each). The IR signals were quantified with an Odyssey infrared scanner (LI-COR Biosciences). The antibodies and concentrations are listed in **Supplementary Table 1**.

#### Quantitative real time-PCR

Total RNA was extracted with the RNeasy kit (Qiagen) and treated with DNase I (Ambion) according to the manufacturer’s recommendation. RNA was reverse-transcribed with random primers using SuperScript III cDNA Synthesis Kit (Invitrogen). cDNA samples were diluted in nuclease-free water and stored at –80°C. RT-PCR reactions were performed on a QuantStudio5 system (ThermoFisher Scientific) with SYBR Green PCR Master Mix (Bioland). Transcript levels were calculated by the comparative cycle threshold method and normalized to cyclophilin A expression. All primers used in the experiments are listed in **Supplementary Table 2**.

#### Doxycycline (Dox)-inducible expression in SMCs

SMCs harboring Dox-inducible pTRIPZ expression vectors for human prelamin A and human progerin have been described previously (*15, 21*). All plasmids were verified by DNA sequencing. Packaging of lentivirus and transduction of cells were performed by UCLA’s Vector Core. Transduced cells were selected with 3 µg/ml puromycin for two weeks; individual clones were isolated by limiting dilution in 96-well plates. Clones were screened by western blotting and immunofluorescence staining. A minimum of 2 clones were isolated for each cell line.

#### Constitutive expression of nuclear localized GFP and lamins in SMCs

SMCs expressing green fluorescent protein (GFP) with a nuclear localization signal (nls-GFP) have been described previously (*21, 31*). Constitutive expression of human versions of prelamin A and progerin in SMCs was performed by transducing SMCs with pCLNR (#17735; Addgene) retroviral expression plasmids. The prelamin A and progerin cDNAs were subcloned into *Hind*III and *Not*I sites of pCLNR by In-Fusion cloning (Takara Bio). Human prelamin A cDNA was amplified with forward primer 5′-GCTAGCGAATTATGGAGACCCCGTCCCAGC-3′ and reverse primer 5′-GATCCTTGCGGCCTTACATGATGCTGCAGT-3′; human progerin cDNA was amplified with forward primer 5′-GCTAGCGAATTATGGAGACCCCGTCCCAGC-3′ and reverse primer 5′-CAGATCCTTGCGGCCTTACATGATGCTGCA-3′. All plasmid DNAs were prepared with Maxiprep kit (Qiagen) and verified by DNA sequencing. Packaging of virus and viral transduction were performed by UCLA’s Vector Core. Transduced cells were selected with 3 µg/ml blasticidin or puromycin for two weeks; individual clones were isolated by limiting dilution. A minimum of 2 clones were isolated for each cell line.

---

**Fig. S1.**
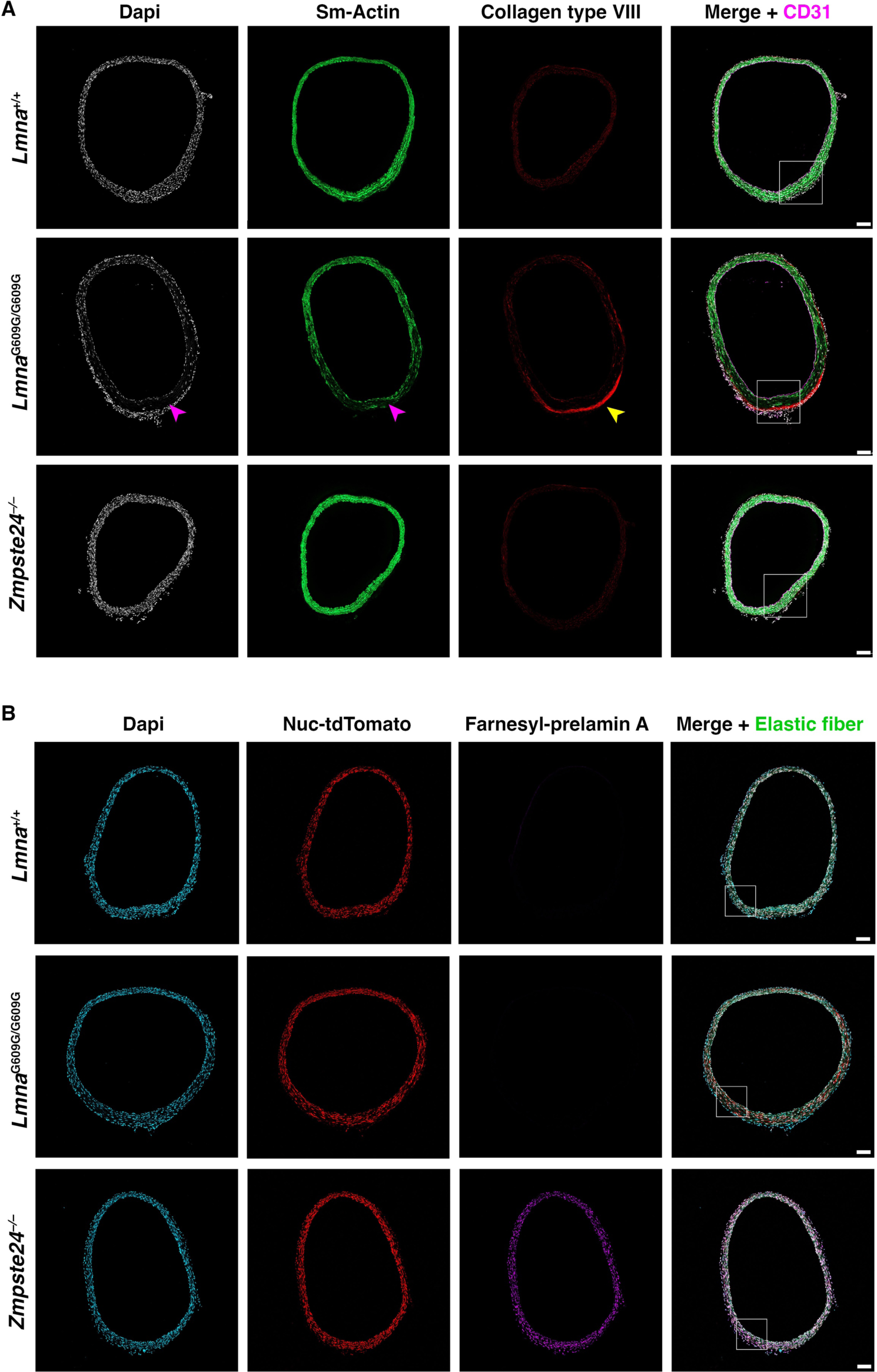
SMC loss and nuclear membrane ruptures are absent in the aorta of *Zmpste24*^−/–^ mice. **A.** Microscopy images of the proximal ascending aorta from 16-week-old *Lmna*^+/+^, *Lmna*^G609G/G609G^, and *Zmpste24*^−/–^ mice stained with antibodies against smooth muscle actin (Sm-actin, *green*), collagen type VIII (*red*), and CD31 (*magenta*). Nuclei were stained with Dapi (*white*). *Red* arrowheads point to areas with reduced numbers of SMC nuclei and reduced Sm-actin staining. The *yellow* arrowhead points to collagen type VIII staining in the adventitia. Scale bar, 100 µm. The boxed regions in the merged images are shown at higher magnification in Fig. 1A. **B**. Microscopy images of the proximal ascending aorta from 13-week-old *Lmna*^+/+^ and *Lmna*^G609G/G609G^ mice and a 21-week-old *Zmpste24*^−/–^ mouse [all expressing a nuclear-targeted tdTomato (Nuc-tdTomato) transgene] stained with an antibody against farnesyl-prelamin A. The images show Dapi (*blue*), Nuc-tdTomato (*red*), farnesyl-prelamin A (*magenta*), and elastic fibers (*green*) in the merged image. Scale bar, 100 µm. The boxed regions in the merged images are shown at higher magnification in Fig. 1B.

**Fig. S2.**
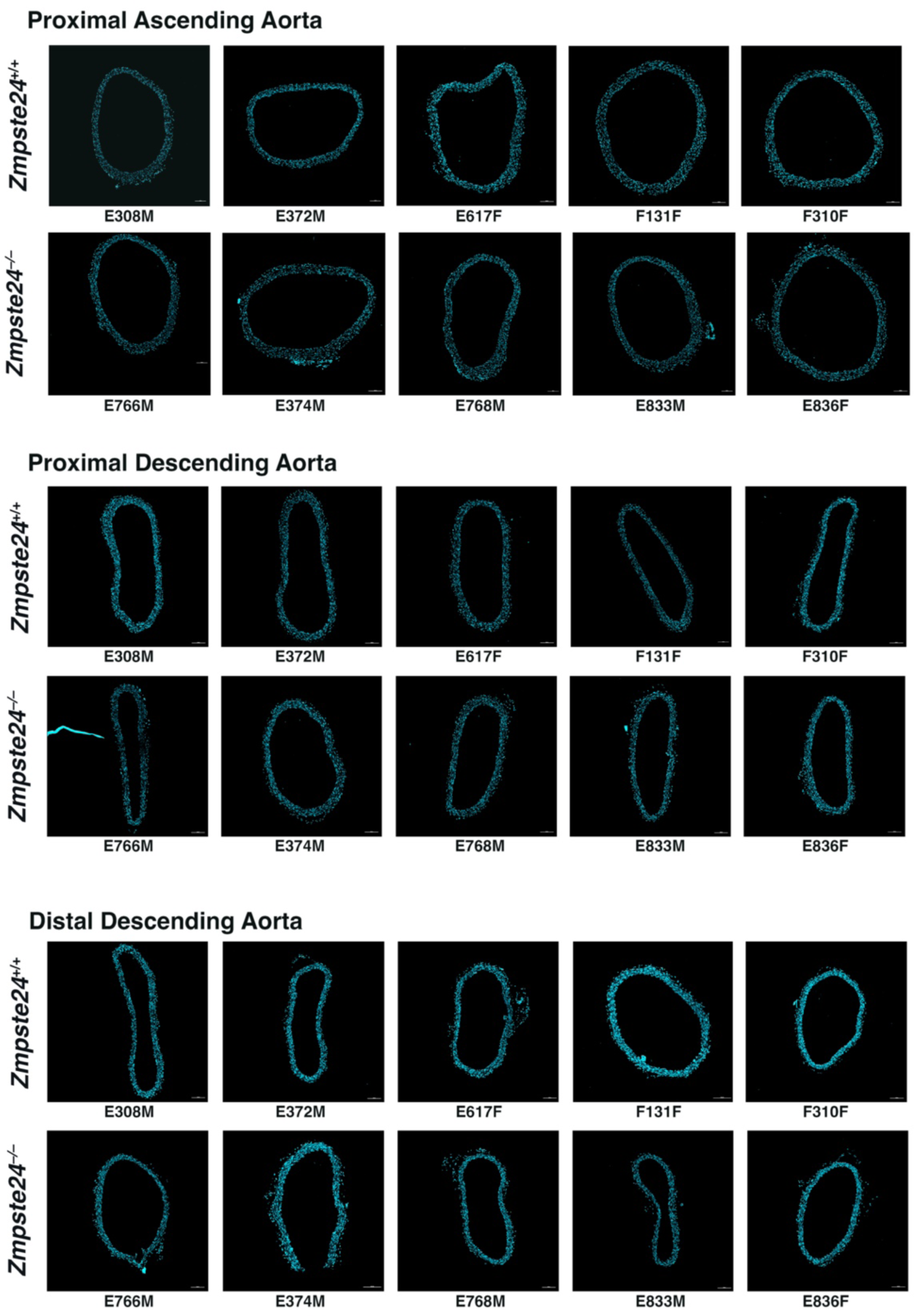
Loss of smooth muscle cells is not detected in the thoracic aorta of *Zmpste24*^−/–^ mice. Microscopy images of the proximal ascending, proximal descending, and distal descending aorta from 21-week-old *Zmpste24*^+/+^ and *Zmpste24*^−/–^ mice (5 mice/group) stained with Dapi (*blue*). The mouse IDs are shown below each image. Scale bar, 100 µm. The images of the proximal ascending aorta were used to generate the quantitative data reported in Fig. 1F.

**Fig. S3.**
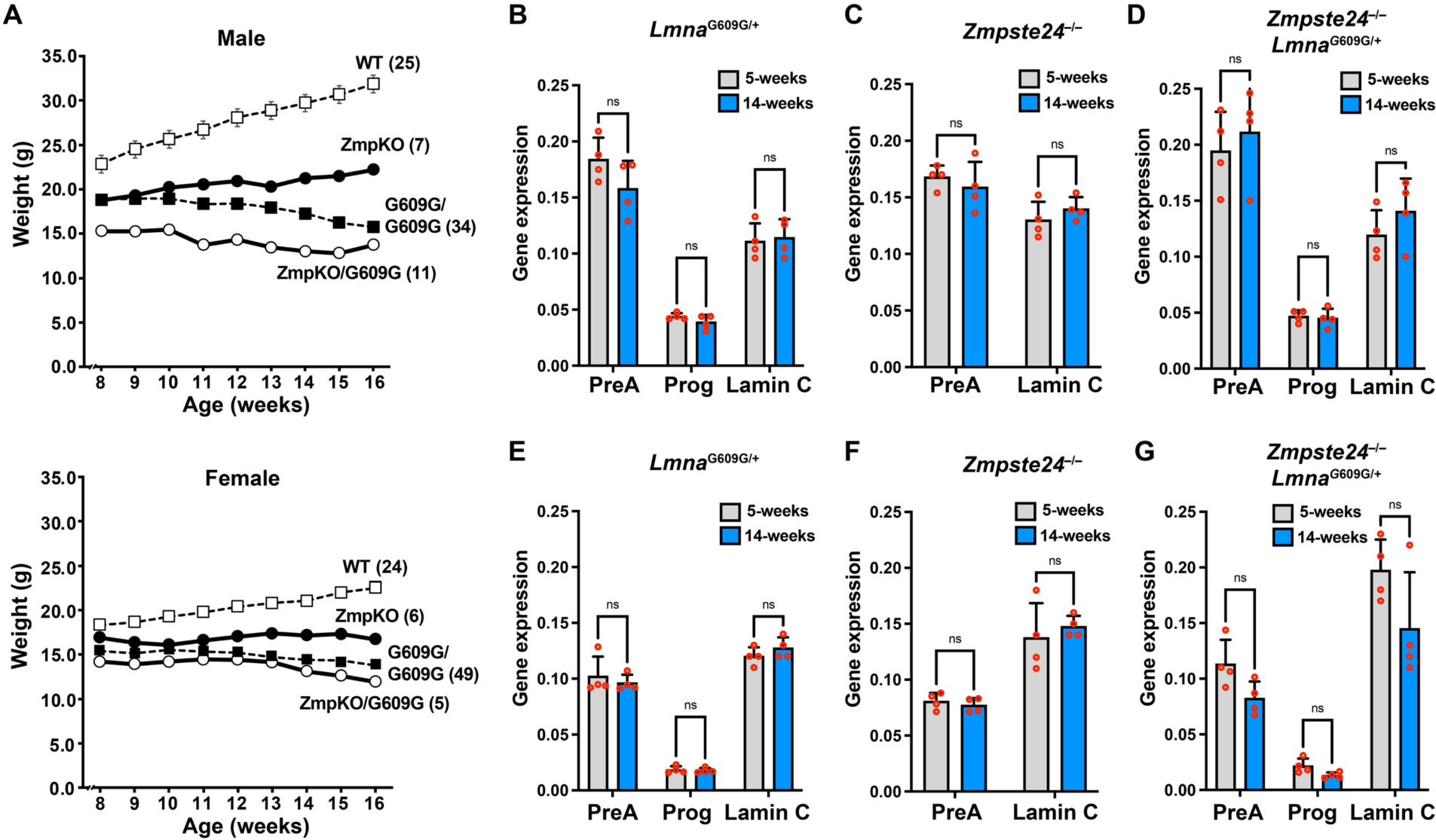
Transcript levels for the A-type nuclear lamins do not increase with age in the aorta or heart of *Lmna*^G609G/+^, *Zmpste24*^−/–^, or *Zmpste24*^−/–^*Lmna*^G609G/+^ mice. **A**. Body weight curves for male (upper) and female (lower) *Lmna*^+/+^ (WT), *Lmna*^G609G/G609G^ (G609G/G609G), *Zmpste24*^− /–^ (ZmpKO), and *Zmpste24*^−/–^*Lmna*^G609G/+^ (ZmpKO/G609G) mice. The weight curves for the *Lmna*^+/+^ and *Lmna*^G609G/G609G^ mice (dotted lines) are from animals generated in a different mouse colony. The numbers of mice per group are shown in parentheses. Mean ± SEM. The error bars for some data points are too small to see. **B–D.** Bar graphs showing prelamin A (PreA), progerin (Prog), and lamin C gene expression (relative to *Ppia*) in aortas from young and old *Lmna*^G609G/+^, *Zmpste24*^−/–^, and *Zmpste24*^−/–^*Lmna*^G609G/+^ mice. Mean ± SEM (*n* = 4 mice/group). Student’s *t* test. ns, not significant. **E–G.** Bar graphs showing PreA, Prog, and lamin C gene expression (relative to *Ppia*) in hearts from young and old *Lmna*^G609G/+^, *Zmpste24*^−/–^, and *Zmpste24*^−/–^*Lmna*^G609G/+^ mice. Mean ± SEM (*n* = 4 mice/group). Student’s *t* test. ns, not significant.

**Fig. S4.**
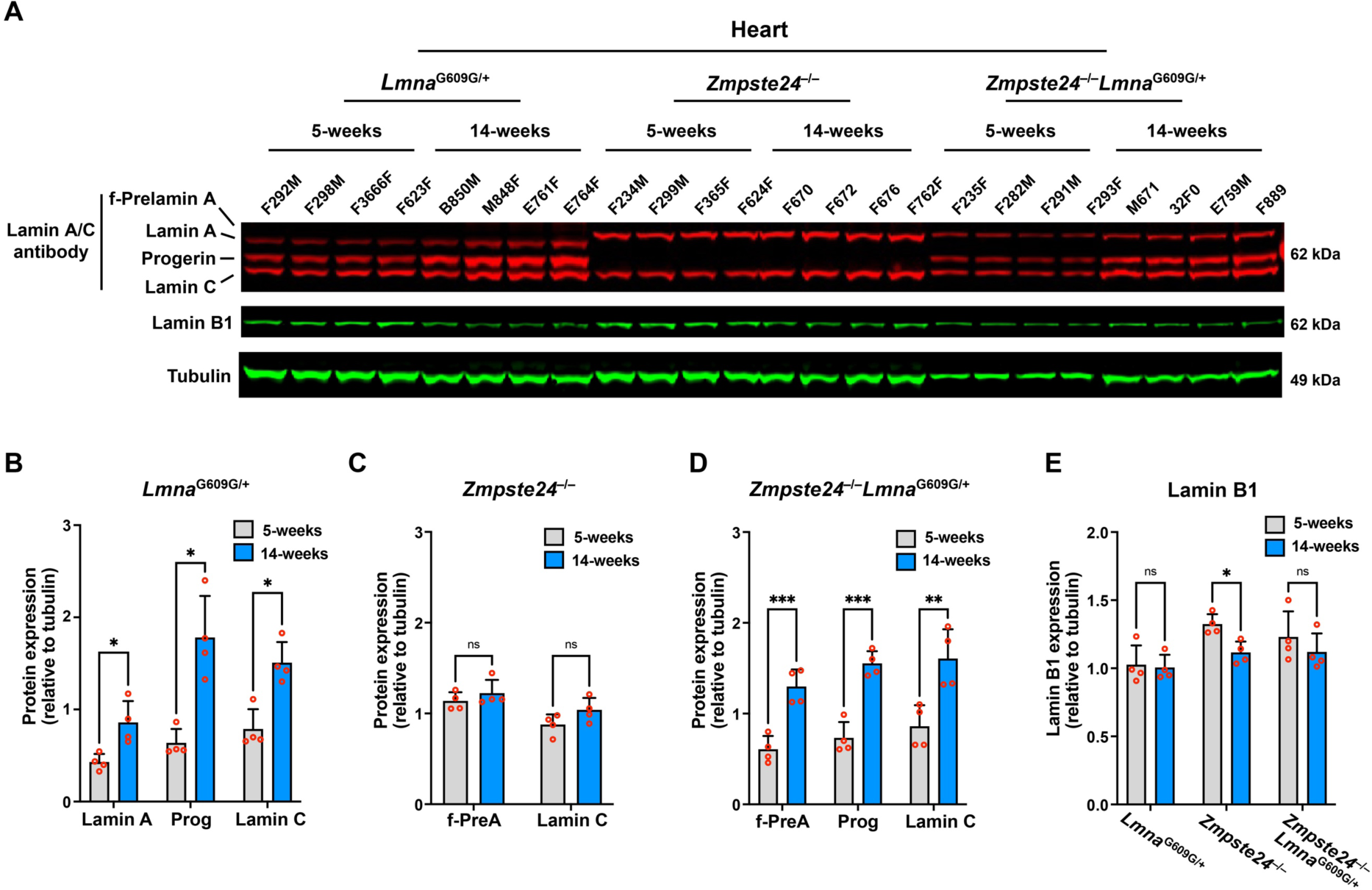
Progerin causes the accumulation of the A-type nuclear lamins in the heart. **A.** Western blot comparing the expression of lamin A, lamin C, farnesyl-prelamin A, progerin, and lamin B1 in hearts from 5- and 14-week-old *Lmna*^G609G/+^, *Zmpste24*^−/–^, and *Zmpste24*^−/–^*Lmna*^G609G/+^ mice. Tubulin was measured as a loading control. The ages and mouse IDs are shown above each sample. **B**. Bar graph showing lamin A, progerin (Prog), and lamin C expression (relative to tubulin) in hearts from young and old *Lmna*^G609G/+^ mice. Mean ± SEM (*n* = 4 mice/group). Student’s *t* test. *, *P* < 0.05. **C**. Bar graph showing farnesyl-prelamin A (f-PreA) and lamin C expression (relative to tubulin) in hearts from young and old *Zmpste24*^−/–^ mice. Mean ± SEM (*n* = 4 mice/group). Student’s *t* test. ns, not significant. **D.** Bar graph showing f-PreA, Prog, and lamin C expression (relative to tubulin) in hearts from young and old *Zmpste24*^−/–^*Lmna*^G609G/+^ mice. Mean ± SEM (*n* = 4 mice/group). Student’s *t* test. **, *P* < 0.01. ***, *P* < 0.001. **E.** Bar graph showing lamin B1 expression (relative to tubulin) in aortas from young and old *Lmna*^G609G/+^, *Zmpste24*^−/–^, and *Zmpste24*^−/–^*Lmna*^G609G/+^ mice. Mean ± SEM (*n* = 4 mice/group). Student’s *t* test. *, *P* < 0.05. ns, not significant.

**Fig. S5.**
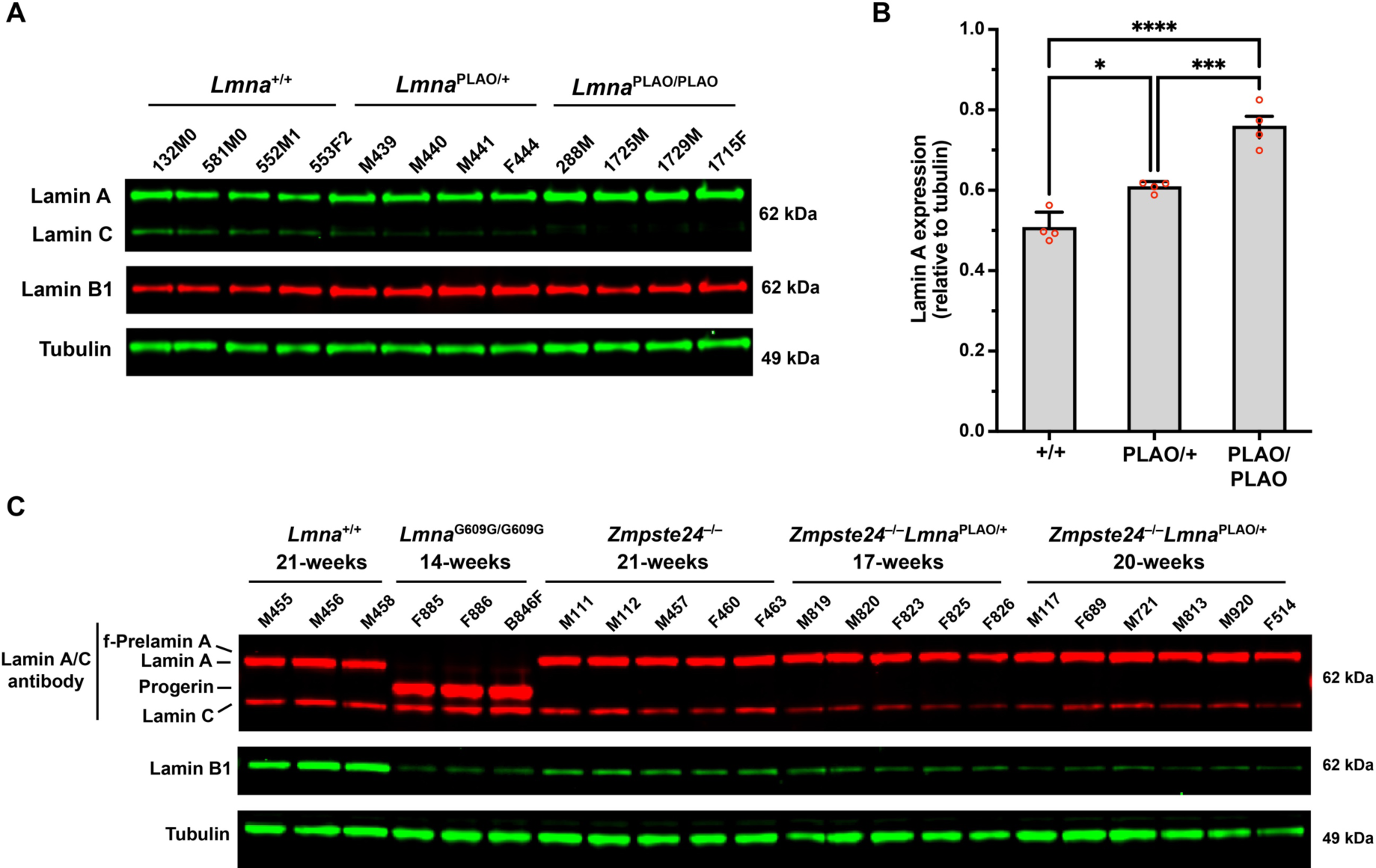
The *Lmna*^PLAO^ allele increases lamin A levels in the aorta of *Zmpste24*^+/+^ mice but does not increase farnesyl-prelamin A levels in *Zmpste24*^−/–^ mice. **A.** Western blot comparing the expression of lamin A, lamin C, and lamin B1 in aortas from *Lmna*^+/+^, *Lmna*^PLAO/+^, and *Lmna*^PLAO/PLAO^ mice. Tubulin was measured as a loading control. The mouse IDs are shown above each sample. **B**. Bar graph showing lamin A expression (relative to tubulin) in aortas from *Lmna*^+/+^ (+/+), *Lmna*^PLAO/+^ (PLAO/+), and *Lmna*^PLAO/PLAO^ (PLAO/PLAO) mice. Mean ± SEM (*n* = 4 mice/group). ANOVA. *, *P* < 0.05. ***, *P* < 0.001. ****, *P* < 0.0001. **C**. Western blot comparing the expression of lamin A, lamin C, progerin, farnesyl-prelamin A, and lamin B1 in aortas from *Lmna*^+/+^, *Lmna*^G609G/G609G^, *Zmpste24*^−/–^, and *Zmpste24*^−/–^*Lmna*^PLAO/+^ mice. Tubulin was measured as a loading control. The mouse IDs and ages are shown above each sample. The levels of farnesyl-prelamin A in 21-week-old *Zmpste24*^−/–^ mice (*n* = 5) and 20-week-old *Zmpste24*^−/–^*Lmna*^PLAO/+^ mice (*n* = 6) were judged not to be different (ANOVA; P = 0.12).

**Fig. S6.**
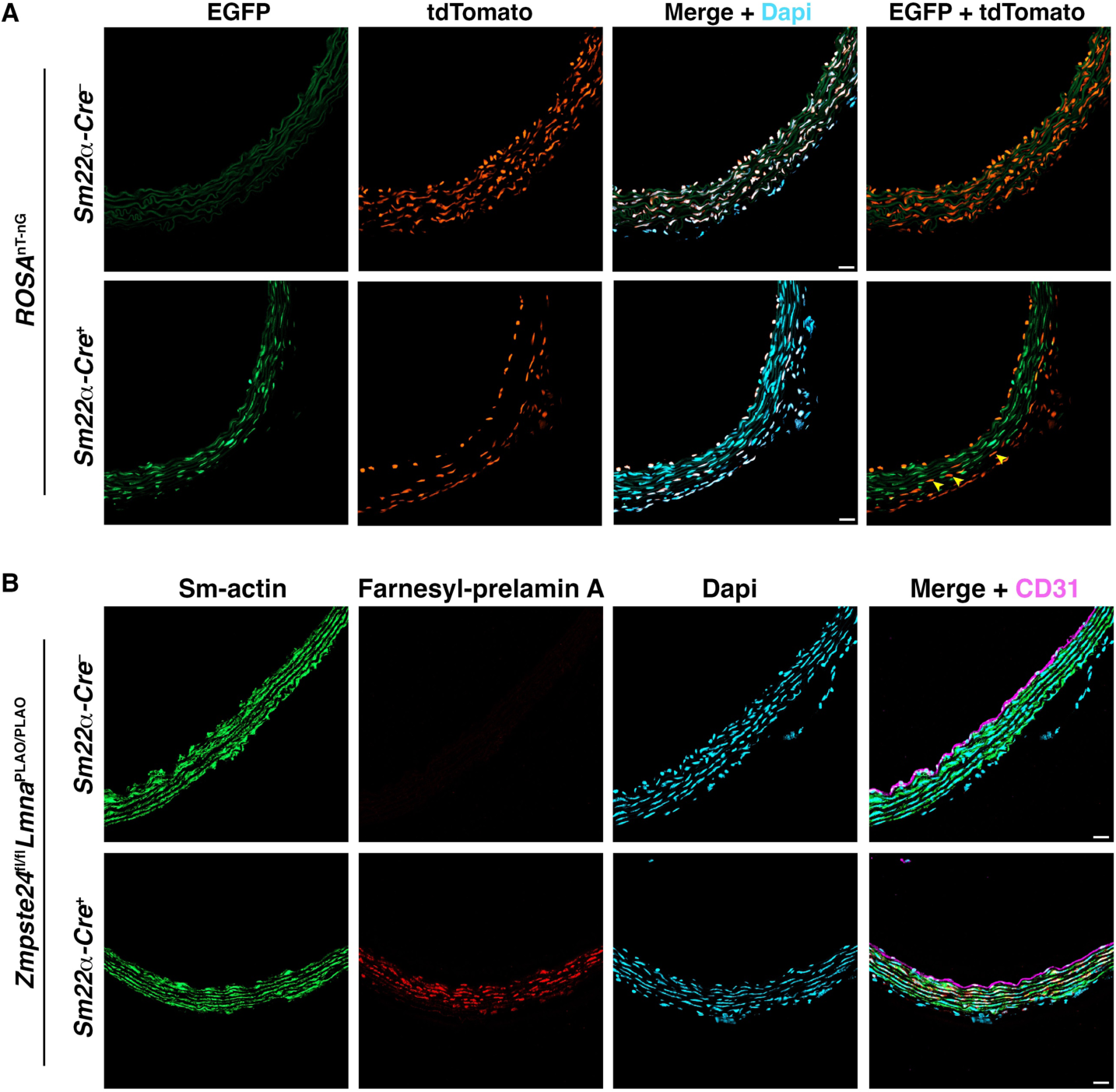
Inactivation of *Zmpste24* expression in SMCs results in farnesyl-prelamin A synthesis in aortic SMCs. **A.** Confocal fluorescence microscopy images of the proximal ascending aorta of an 8-week-old *Sm22α-Cre*^−^*Rosa*^nT-nG^ (upper row) and *Sm22α-Cre*^+^*Rosa*^nT-nG^ (lower row) mouse. The *Rosa*^nT-nG^ allele is a two-color reporter that expresses tdTomato in the nucleus (*39*). After *Cre* recombination, the *Rosa*^nT-nG^ allele expresses EGFP, identifying cells that express *Cre*. The images show EGFP (*green*), tdTomato (*red*), and Dapi (*blue*). The *yellow* arrowheads point to SMCs in an *Sm22α-Cre*^+^*Rosa*^nT-nG^ mouse that still express tdTomato, showing that *Cre* is not expressed in all aortic SMCs of *Sm22α-Cre*^+^ mice. Scale bar, 20 µm. **B.** Confocal fluorescence microscopy images of cryosections from the proximal ascending aorta of a 4-week-old *Sm22α-Cre*^−^*Zmpste24*^fl/fl^*Lmna*^PLAO/PLAO^ mouse (upper row) and *Sm22α-Cre*^+^*Zmpste24*^fl/fl^*Lmna*^PLAO/PLAO^ mouse (lower row) stained with antibodies against smooth muscle actin (Sm-actin, *green*), farnesyl-prelamin A (*red*), and CD31 (*magenta*). Nuclei were stained with Dapi (*blue*). Scale bar, 20 µm.

**Fig. S7.**
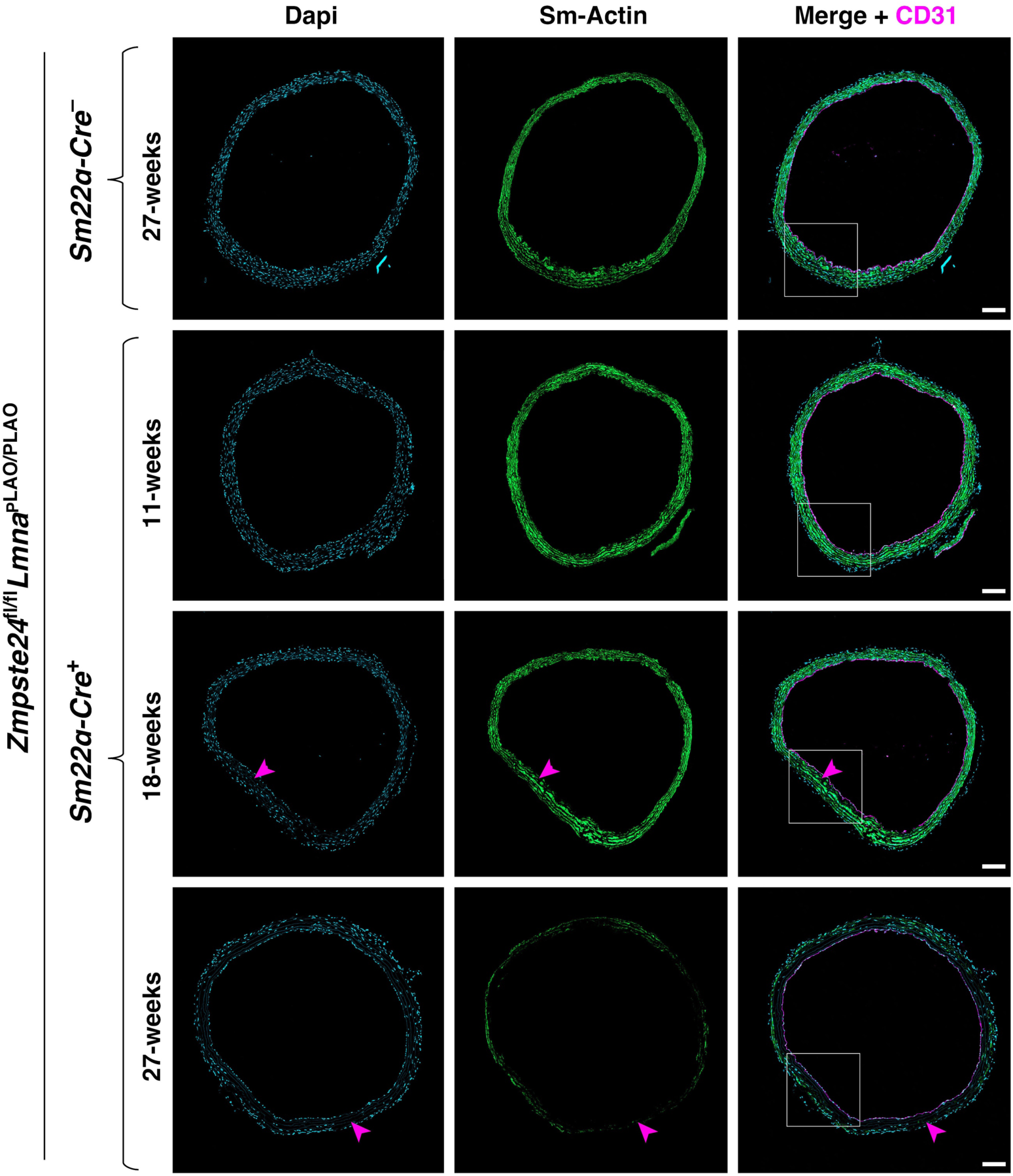
SMC loss and reduced smooth muscle actin staining in *Sm22α-Cre*^+^*Zmpste24*^fl/fl^*Lmna*^PLAO/PLAO^ mice. Confocal fluorescence microscopy images of the proximal ascending aorta from 11-, 18-, and 27-week-old *Sm22α-Cre^+^Zmpste24*^fl/fl^*Lmna*^PLAO/PLAO^ mice stained with antibodies against smooth muscle actin (Sm-actin, *green*) and CD31 (*magenta*). Nuclei were stained with Dapi (*blue*). As a control, images from a 27-week-old *Sm22α-Cre*^−^ *Zmpste24*^fl/fl^*Lmna*^PLAO/PLAO^ mouse are shown. Scale bar, 100 µm. *Red* arrowheads point to areas with reduced Sm-actin staining and reduced numbers of SMC nuclei. The boxed regions are shown at higher magnification in Fig. 5C.

**Fig. S8.**
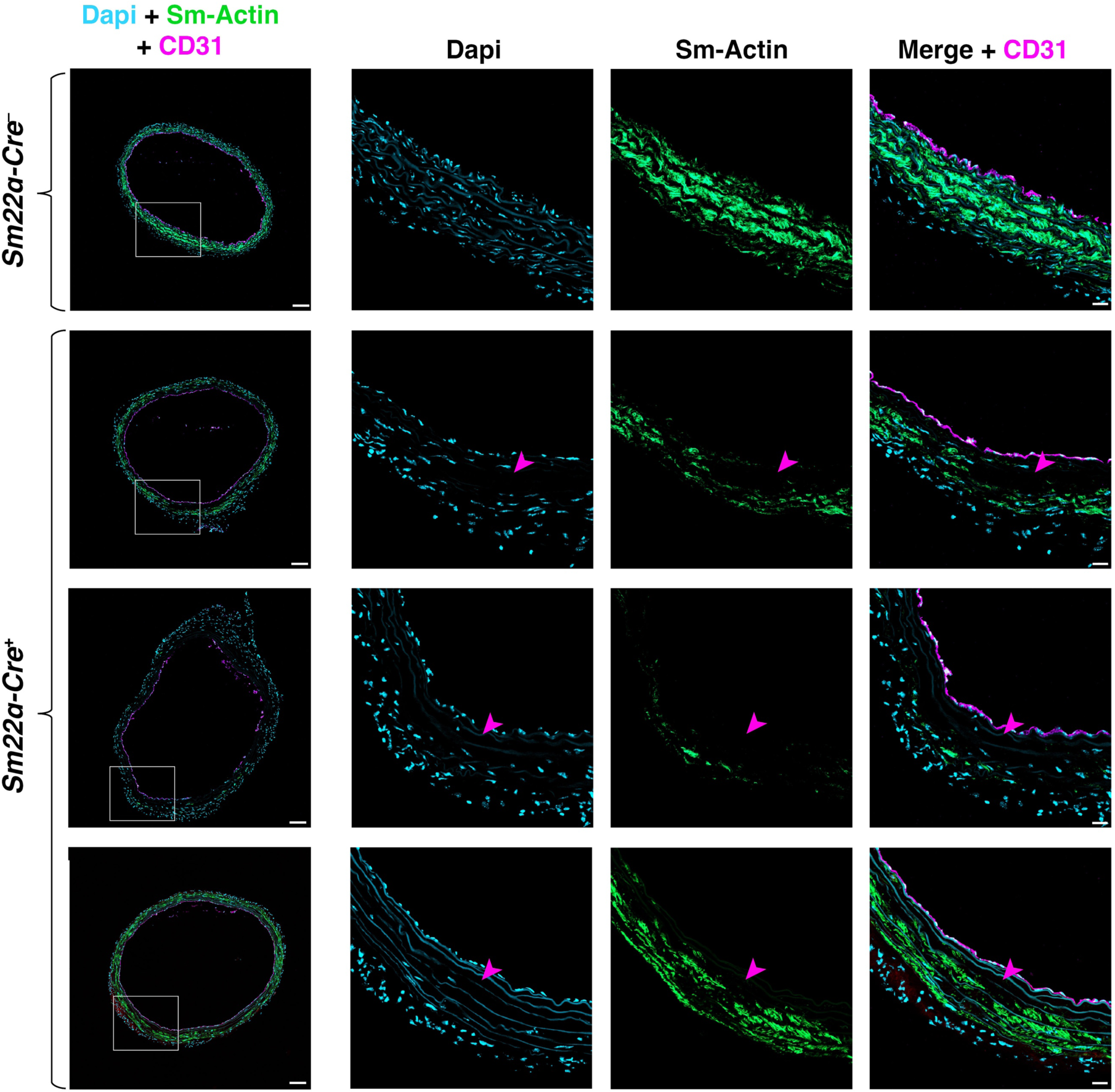
Farnesyl-prelamin A causes SMC loss in the aorta of 27-week-old *Sm22α-Cre*^+^*Zmpste24*^fl/fl^*Lmna*^PLAO/PLAO^ mice. Confocal fluorescence microscopy images of the proximal ascending aorta from 27-week-old *Sm22α-Cre^+^Zmpste24*^fl/fl^*Lmna*^PLAO/PLAO^ mice stained with antibodies against smooth muscle actin (Sm-actin, *green*) and CD31 (*magenta*). Nuclei were stained with Dapi (*blue*). As a control, images from a 27-week-old *Sm22α-Cre*^−^ *Zmpste24*^fl/fl^*Lmna*^PLAO/PLAO^ mouse are shown. Scale bar, 100 µm. Each row represents a different mouse. The boxed regions are shown at higher magnification to the right. *Red* arrowheads point to areas with reduced Sm-actin staining and reduced numbers of SMC nuclei. Scale bar, 20 µm.

**Table S1.** Antibodies used for western blotting and immunocytochemistry.

| **Antibody Description** | **Species** | **Source** | **Catalog #** | **Use** | **Dilution** |
| --- | --- | --- | --- | --- | --- |
| Akt (pan) | Rabbit | Cell Signaling | 4691 | WB | 1:1000 |
| β-actin | Mouse | Santa Cruz Biotech | SC47778 | WB | 1:3000 |
| CD31 (clone 2H8) | Hamster | DSHB | AB_2161039 | IF | 10 μg/ml |
| Collagen Type VIII | Rabbit | Antibodies-online | ABIN1718654 | IF | 1:20 |
| Human lamin A | Mouse | Millipore | MAB3211 | WB,<br>IF | 1:1500 |
| Human lamin A/C | Rabbit | Abcam | Ab108595 | WB,<br>IF | 1:1000 |
| Human lamin A/C-Alexa 488 conjugated | Rabbit | Abcam | Ab185014 | IF | 1:1000 |
| Lamin A/C | Mouse | Santa Cruz Biotech | SC376248 | WB,<br>IF | 1:1000 |
| Lamin B1 | Goat | Santa Cruz Biotech | SC6217 | WB,<br>IF | 1:1500 |
| Lamin B1 | Rabbit | Invitrogen | 702972 | WB,<br>IF | 1:1000 |
| Prelamin A | Rat | In-house | clone 3C8 | WB | 1:1000 |
| Phospho-Lamin A/C (Ser404) | Rabbit | Millipore Sigma | ABT1387 | WB | 1:500 |
| Phospho-Akt (Ser473) | Rabbit | Cell Signaling | 4060 | WB | 1:1000 |
| Sm-actin | Goat | Sigma | SAB2500963 | IF | 1:100 |
| Tubulin | Rat | Novus Bio | NB600-506 | WB | 1:3000 |
| Anti-rabbit IR800 | Donkey | LI-COR | 926-32213 | WB | 1:10000 |
| Anti-goat IR800 | Donkey | LI-COR | 926-32214 | WB | 1:10000 |
| Anti-rat IR800 | Donkey | ThermoFisher | SA5-10032 | WB | 1:5000 |
| Anti-mouse IR800 | Donkey | ThermoFisher | SA5-10172 | WB | 1:5000 |
| Anti-rabbit IR680 | Donkey | LI-COR | 926-32221 | WB | 1:5000 |
| Anti-rat IR680 | Goat | LI-COR | 925-68076 | WB | 1:5000 |
| Anti-goat IR680 | Donkey | LI-COR | 926-68074 | WB | 1:5000 |
| Anti-mouse IR680 | Donkey | ThermoFisher | SA5-10170 | WB | 1:5000 |
| Anti-mouse Alexa 488 | Donkey | Invitrogen | A21202 | IF | 1:2000 |
| Anti-rabbit Alexa 488 | Donkey | Invitrogen | A21206 | IF | 1:2000 |
| Anti-goat Alexa 488 | Donkey | Invitrogen | A11055 | IF | 1:2000 |
| Anti-goat Alexa 555 | Donkey | Invitrogen | A21432 | IF | 1:2000 |
| Anti-rabbit Alexa 555 | Donkey | Invitrogen | A31572 | IF | 1:200 |
| Anti-rabbit Alexa 568 | Donkey | Invitrogen | A10042 | IF | 1:2000 |
| Anti-mouse Alexa 568 | Donkey | Invitrogen | A10037 | IF | 1:2000 |
| Anti-mouse Alexa 647 | Donkey | Invitrogen | A31571 | IF | 1:2000 |

**Table S2.**
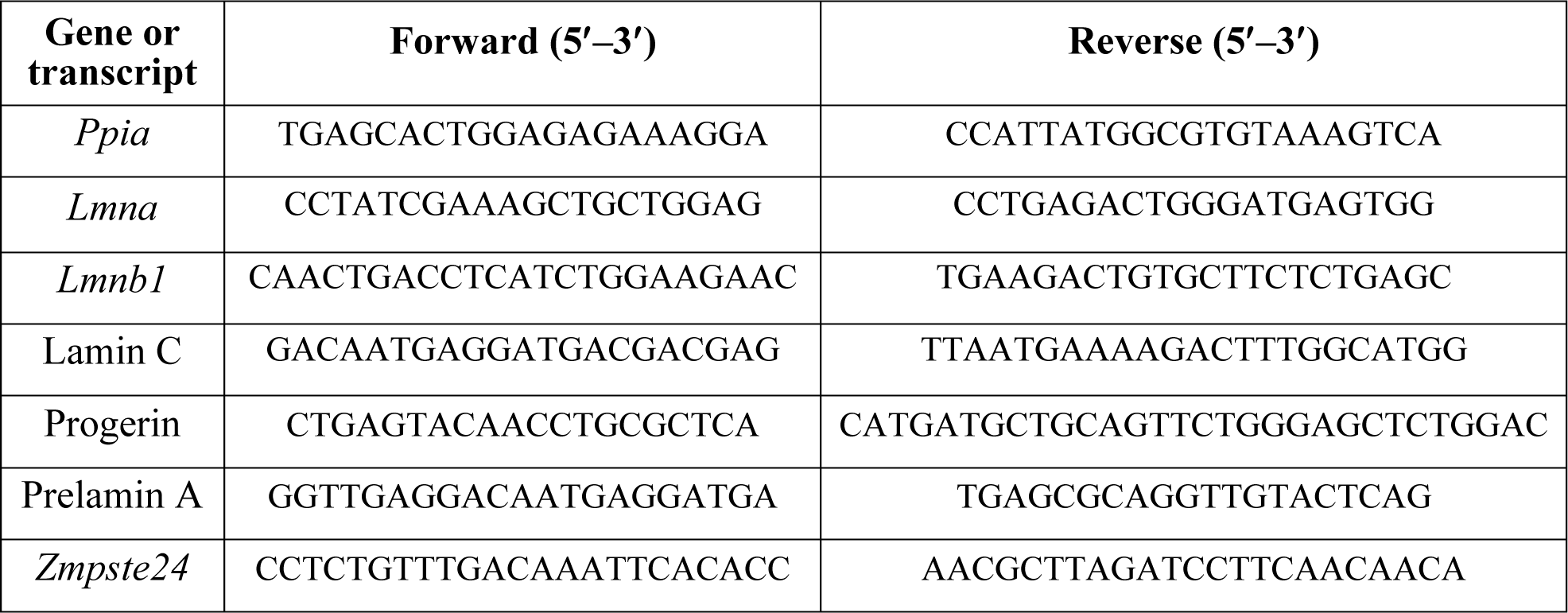
Quantitative RT-PCR primers.

**Table S3.** Description of cell lines.

| Cell line | Modification | Source | Validation method | Mycoplasma contamination |
| --- | --- | --- | --- | --- |
| hu-prelamin A-SMC | human (hu)-prelamin A-pTRIPZ | In-house | Western blotting and microscopy | No |
| hu-progerin-SMC | hu-progerin-pTRIPZ | In-house | Western blotting and microscopy | No |
| hu-prelamin A-SMC + nls-GFP | hu-prelamin A-pTRIPZ + nls-GFP-pCLNR | In-house | Microscopy | No |
| hu-progerin-SMC + nls-GFP | hu-progerin-pTRIPZ + nls-GFP-pCLNR | In-house | Microscopy | No |
| *Zmpste24*<sup>-/-</sup>-SMC | CRISPR/Cas9 deletion of *Zmpste24* | In-house | Western blotting, qPCR and sequencing | No |
| *Lmna*<sup>-/-</sup>-SMC | CRISPR/Cas9 deletion of *Lmna* | In-house | Western blotting, qPCR and sequencing | No |
| *Zmpste24*<sup>-/-</sup>-SMC + hu-prelamin A-pTRIPZ | human (hu)-prelamin A-pTRIPZ | In-house | Western blotting and microscopy | No |
| *Zmpste24*<sup>-/-</sup>-SMC + nls-GFP | *Zmpste24*<sup>-/-</sup> + nls-GFP-pCLNR | In-house | Microscopy | No |
| *Zmpste24*<sup>-/-</sup>-SMC + hu-progerin | *Zmpste24*<sup>-/-</sup> + hu-progerin-pCLNR | In-house | Western blotting and microscopy | No |

